# DNA oxidation induced by fetal exposure to BPA agonists impairs female meiosis

**DOI:** 10.1101/2020.08.17.253724

**Authors:** Sonia Abdallah, Delphine Moison, Margaux Wieckowski, Sébastien Messiaen, Emmanuelle Martini, Anna Campalans, J. Pablo Radicella, René Habert, Gabriel Livera, Virginie Rouiller-Fabre, Marie-Justine Guerquin

## Abstract

Many endocrine disruptors have been proven to impair the meiotic process that is mandatory to produce healthy gametes. Bisphenol A is emblematic as it impairs meiotic prophase I and causes oocyte aneuploidy following *in utero* exposure. However, the mechanisms underlying these deleterious effects remain poorly understood. Furthermore, the increasing uses of BPA analogs raise concerns for public health. Here, we investigated the effect on oogenesis in mouse of fetal exposure to two BPA analogs, Bisphenol A Diglycidyl Ether (BADGE) or Bisphenol AF (BPAF). These analogs delay meiosis initiation, increase MLH1 foci *per* cell and induce oocyte aneuploidy. We further demonstrate that these defects are accompanied by a deregulation of gene expression and aberrant mRNA splicing in fetal premeiotic germ cells. Interestingly, we observed an increase in DNA oxidation after exposure to BPA analogs. Specific induction of oxidative DNA damages during fetal germ cell differentiation causes similar defects during oogenesis, as observed in 8-Oxoguanine DNA Glycosylase (OGG1) deficient mice or after *in utero* exposure to potassium bromate (KBrO3), an inducer of oxidative DNA damages. Moreover, the supplementation of N-acetylcysteine (NAC) with BPA analogs counteracts the bisphenol-induced meiotic effect. Together our results position oxidative stress as a central event that negatively impacts the female meiosis with major consequences on oocyte quality. This could be a common mechanism of action for so called endocrine disruptors pollutants and it could lead to novel strategies for reprotoxic compounds.

## Introduction

In females, aneuploidy (aberrant number of chromosomes) is an important cause of adverse reproductive outcomes such as miscarriages and congenital abnormalities. Aneuploid eggs can be induced by numerous lifestyle factors (age, obesity, environmental pollutants…) (Nagaoka et al., 2012) and can derive from alterations occurring during meiotic prophase I in fetal life. Indeed, proper chromosome segregation at adulthood requires organized reciprocal DNA exchanges between homologous chromosomes (crossover) occurring during prophase I as a result of the repair of meiotic double strand break (DSB) by homologous recombination (Patricia A. Hunt & Hassold, 2008; Ottolini et al., 2015; S. Wang et al., 2017). Crossover regulation depends on a correct implementation of the meiotic program in primordial germ cells (PGCs) initiated after their migration into the gonad. At this stage, pluripotent and proliferative PGCs acquire the competence to initiate meiosis through the expression of Deleted In Azoospermia-like (Dazl) (Nicholls et al., 2019). Depending on the somatic environment and under the control of meiotic orchestrators such as *Stra8* that direct the switch from mitosis to meiosis, female PGCs initiate prophase I at 13.5 day post-conception (dpc) (Bailey et al., 2017; Hargan-Calvopina et al., 2016a; Ishiguro et al., 2020; Le Bouffant et al., 2010; Spiller & Bowles, 2019; Trautmann et al., 2008). Alterations that occur during the establishment of the meiotic program lead to meiotic defects in prophase I that can hamper future fertility (Bailey et al., 2017; Hargan-Calvopina et al., 2016a; Ishiguro et al., 2020; Nicholls et al., 2019). It is well known that implementation and progression of prophase I are extremely sensitive to environmental factors such as toxicants and endocrine disrupting chemicals. Among those, bisphenol A (BPA) is the first and the most studied environmental compound known to alter meiosis and folliculogenesis in females in numerous mammalian and non-mammalian organisms (Brieno-Enriquez et al., 2012; Brieño-Enríquez et al., 2011; P. A. Hunt et al., 2012; Lawson et al., 2011; Susiarjo et al., 2007; W. Wang et al., 2014; H.-Q. Zhang et al., 2012; T. Zhang et al., 2014). In female primates and rodents, fetal exposure to BPA induces alteration of the expression of meiotic genes at the time of meiosis onset and during prophase I (Brieno-Enriquez et al., 2012; Lawson et al., 2011; H.-Q. Zhang et al., 2012). Moreover, fetal exposure to BPA alters the distribution of recombination events signaled by MLH1, a DNA mismatch repair protein required for resolution of double Holliday junctions as crossovers (Ashley et al., 2001; Cheng et al., 2009; Patricia A. Hunt et al., 2003). This increase observed during pachytene stage is correlated with the occurrence of aneuploid oocytes at adulthood. The precise action of BPA on meiosis has been proposed to involve estrogen receptor signaling (Gibert, 2015; Susiarjo et al., 2007; M. Yu et al., 2018; H.-Q. Zhang et al., 2012). Moreover, several studies suggest that BPA exposure induces epigenetic alterations in germ cells such as DNA or histone methylation resulting in gene expression alterations (Chao et al., 2012; Chianese et al., 2017; Kim et al., 2014; T. Wang et al., 2016; H.-Q. Zhang et al., 2012). Lastly, some studies have shown that post-natal exposure to BPA increases the amount of reactive oxygen species (ROS) compromising oocyte maturation and chromosome segregation and inducing DNA damages such as 8-hydroxydeoxyguanosine (8OdG) (Ganesan & Keating, 2016; M. Zhang et al., 2017; T. Zhang et al., 2018).

BPA is a member of the bisphenol family like other structural analogs that are commonly used in the industry. Due to recent regulations and growing commercialization of BPA-free labeled products, BPA analogs are increasingly used in the manufacturing of consumer products. Among those, we choose to focus our study on emerging bisphenols: Bisphenol A Diglycidyl Ether (BADGE) and Bisphenol AF (BPAF) because very little data exists on their effects on mammalian germ cells. However, recent studies have shown that, as BPA, BPAF induces the release of ROS in adult oocytes, delaying *in vitro* maturation (Ding et al., 2017; Jones et al., 2018).

The aim of this study was to explore the effects of prenatal exposure of murine oocytes to BADGE and BPAF and their consequences on fertility at adulthood. We show that exposure of pregnant mice to the BPA analogs, BADGE and BPAF causes oxidative DNA damage, alterations during mitosis to meiosis transition and an increase of crossovers number leading to aneuploidy and oocyte loss. Specific induction of oxidative DNA damages during fetal germ cell differentiation causes similar defects in prophase I as observed in 8-Oxoguanine DNA Glycosylase (OGG1) deficient mice and after fetal exposure to potassium bromate. Thus, we unveiled the central role of oxidative DNA damage in the meiotic response to bisphenols.

## Results

### BADGE or BPAF fetal exposure reproduces BPA defects on folliculogenesis and recombination events

In mammals, fetal exposure to BPA disrupts follicle formation causing multioocyte follicles and increases the incidence of missegregation after meiosis resumption in post-natal ovaries. To study the effects of analogs of BPA on oocyte and folliculogenesis at adulthood, we exposed pregnant mice to 10 µM of BADGE or BPAF in drinking water from 10.5 to 18.5 days post-conception.

At 3 months old, the total number of follicles was assayed in treated (BADGE or BPAF) and control (ethanol [ETOH]) ovaries (Figure 1A-C). A significant decrease of follicular pool was detected in treated ovaries (Figure 1B). The distribution of follicle classes (primordial to antral follicle) was not affected by the treatment suggesting that the progression of folliculogenesis was not affected (Figure 1B). However, we observed abnormal multioocyte follicles in BADGE-and BPAF-treated 3-months ovaries (0 follicle in ETOH group *VS* 34 ± 11 follicles and 58 ± 14 follicles in BADGE and BPAF groups respectively; Figure 1C). The incidence of multioocyte follicles was even more pronounced at the initiation of the folliculogenesis (*ie* 8 days postpartum) in the BADGE- and BPAF-treated ovaries (0 follicle in ETOH group *VS* 647 ± 262 follicles and 75 ± 11 follicles in BADGE and BPAF groups respectively; Figure 1C).

**Figure 1:**
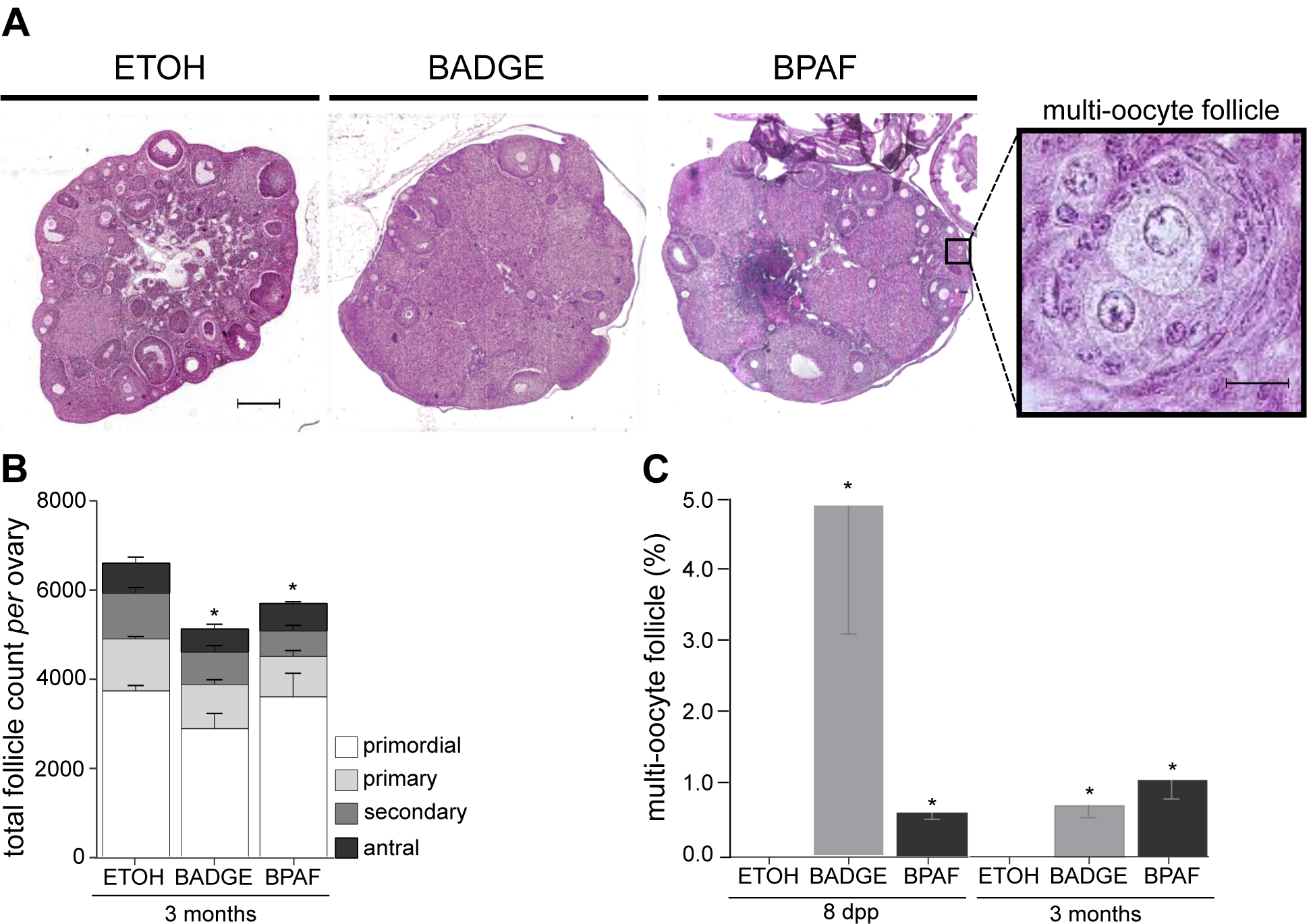
BADGE or BPAF fetal exposures fetal exposure impairs folliculogenesis in post-natal mouse ovaries. (A) Haematoxylin and eosin staining of ovarian section from 3 months old mice exposed during fetal life to vehicle (ETOH), BADGE (BADGE) or BPAF (BPAF). Scale bar, 200µm or 10µm for the higher magnification (multi-oocyte follicle). (B) Quantification of total follicle count in bisphenol-treated or control adult ovaries. Each bar represents the percentage of primordial, primary, secondary and antral follicles. (C) Percentage of multioocytes follicle in bisphenols-treated or control post-natal (8 dpp) and adult (3 months) ovaries. Error bars indicate mean ± s.e.m. n=3-5 mice from 3 independent exposures, * p <0.05 (Mann-Whitney’s test).

As observed with BPA, exposure to bisphenol A analogs induces chromosome missegragation during meiotic divisions. Ploidy in metaphase II oocytes from bisphenols-treated ovaries was assessed by the presence of isolated chromosomes. BADGE or BPAF fetal exposure significantly increases the percentage of aneuploid metaphase II (MII) oocytes (Figure 2A). During prophase I, crossovers and chiasmata distribution in oocyte was also modified after BPAF or BADGE *in utero* exposure. We quantified MLH1 foci in pachytene cells at 18.5 days post-conception (dpc) in bisphenols-treated ovaries. Pachytene stage was identified on the basis of SYCP3 staining patterns and the number of MLH1 foci per bivalent (chromosome pair) was determined. The number of bivalent with three or more MLH1 foci was increased in bisphenols-treated oocytes (Figure 2C). The increase of reciprocal exchanges between homologs was confirmed by the quantification of chiasmata in metaphase I oocytes. In bisphenols-treated oocytes, we observed an increase in bivalent with three or more chiasmata identified according to the shape of the bivalent (Figure 2B). Thus BPAF and BADGE mimic the known hallmarks of BPA during oogenesis like impaired crossovers distribution and aneuploidy.

**Figure 2:**
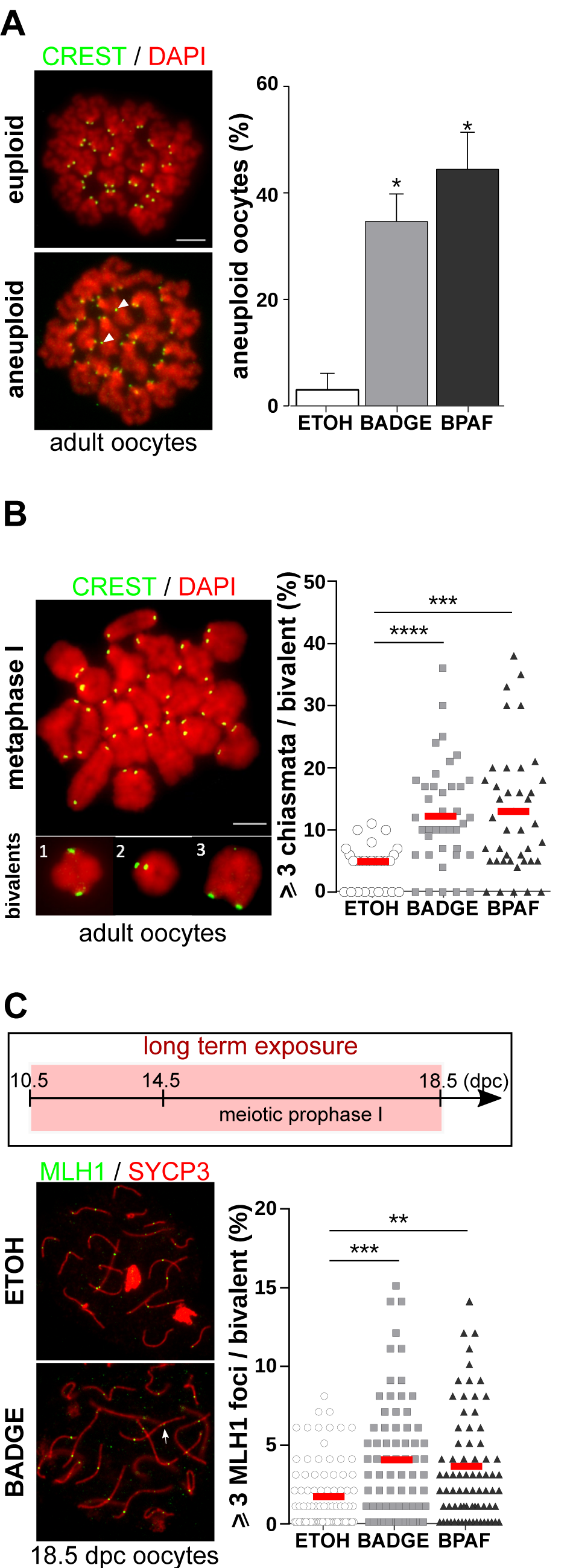
BADGE or BPAF fetal exposures increase the recombination events and missegregation during meiosis. (A) Aneuploid MII oocytes in 3 months old mice exposed during fetal life with vehicle (ETOH), BADGE or BPAF were quantified after CREST immunofluorescence staining. Left panel: photomicrographs of representative euploid (20 chromosomes, 40 centromeres) or aneuploid oocytes (≠20 chromosomes, ≠40 centromeres). Arrowhead indicates extra chromatids. Scale bar: 10 µm. Right panel: quantification of the percentage of aneuploid MII oocytes. n= 28 (ETOH), 40 (BADGE) and 42 (BPAF) oocytes from 5-12 independent exposures. Mean± s.e.m, * p < 0.05 (Mann-Whitney’s test). (B) The number of chiasmata per bivalent (tetrad) in MI oocytes from 3 months old mice exposed during fetal life with vehicle (ETOH), BADGE or BPAF were determined according to the shape of the bivalent after CREST immunostaining (green). Left panel: representative photomicrograph of MI oocyte that contains tetrads with one (1), two (2), three or more (3) chiasmata. Scale bar: 10 µm. Right panel: percentage of bivalent with 3 or more chiasmata. n= 33 (ETOH), 56 (BADGE) and 23 (BPAF) oocytes from 5 independent exposures. Mean (red bar), *** p < 0.001, **** p < 0.0001 (Mann-Whitney’s test). (C) Ovaries from fetuses exposed to vehicle (ETOH) or bisphenols (BADGE and BPAF) from 10.5 dpc to 18.5 dpc were used for the MLH1 quantification (long exposure). The number of crossovers per synaptonemal complexes (synapsis) in pachytene oocytes from 18.5 dpc fetuses was quantified using MLH1 immunostaining (green). The pachytene stage is determined on the basis of the SYCP3 staining (red). Left panel: Representative photomicrographs of pachytene cells from vehicle (ETOH) and BADGE treated ovaries. White arrow shows a synaptonemal complexe with 3 MLH1 foci. Right panel: Percentage of synaptonemal complexes with 3 or more MLH1 foci. n= 86 (ETOH), 78 (BADGE) and 82 (BPAF) oocytes from 12 independent exposures. Mean: red bar, ** *p* < 0.01; \*\*\**p* < 0.001.

### BADGE or BPAF fetal exposures delay meiotic progression and initiation

At 18.5 dpc, we observed a delay in meiotic progression in bisphenols-treated mice with a decrease in the proportion of diplotene stages in favor to the early and late pachytene stages (Figure 3A).

**Figure 3:**
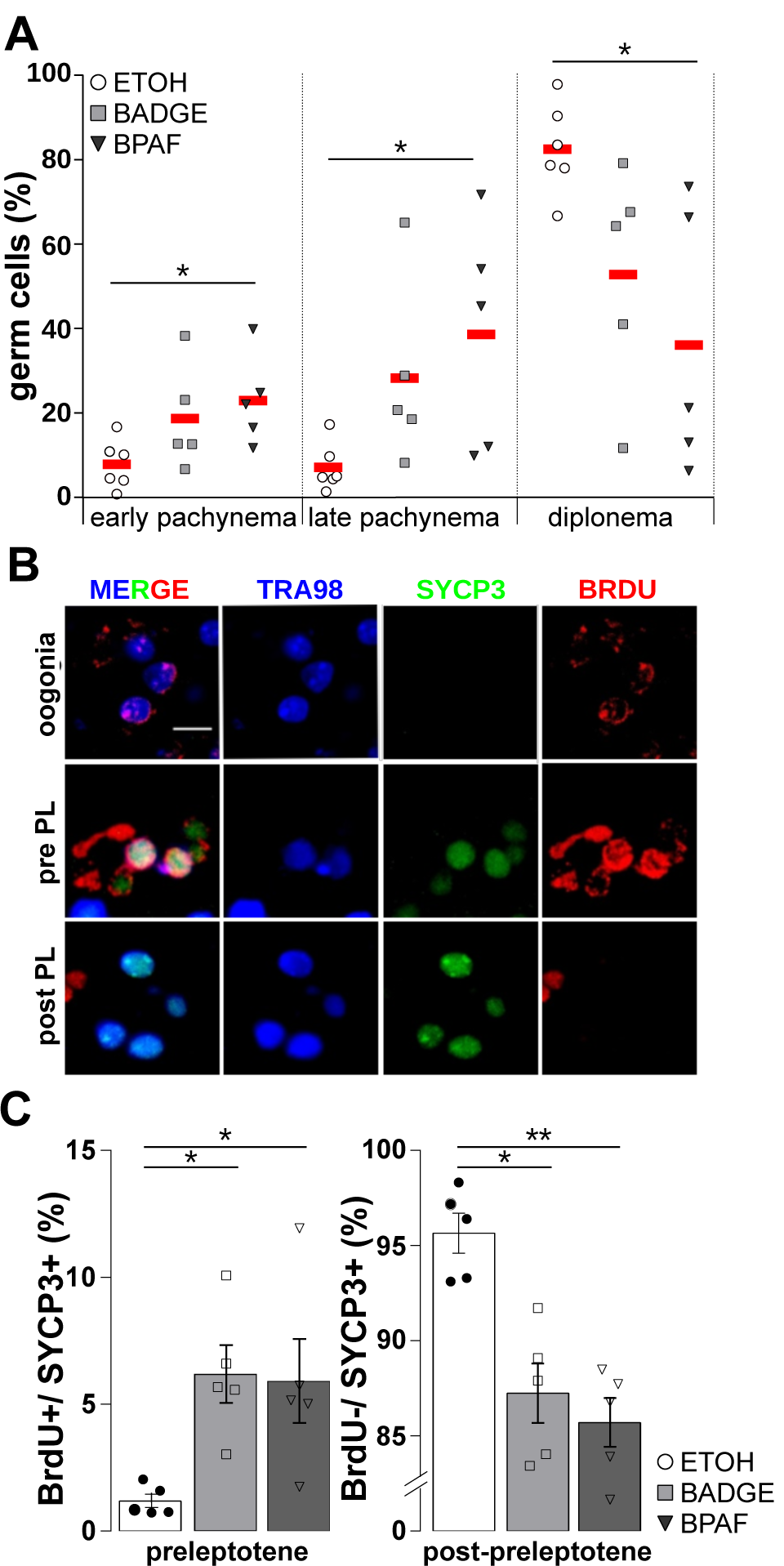
BADGE or BPAF fetal exposures delay the meiosis initiation and prophase I progression. (A) Distribution of meiosis prophase I late stages in 18.5 dpc treated (BADGE, BPAF) and control (ETOH) ovaries. Error bars show mean ± s.e.m.; mice analysed n=3-6; * *p* < 0.05 (Mann-Whitney’s test). (B-C) prophase I initiation was assayed in 14.5 dpc ovaries after immunostaining of SYCP3 (meiosis), BrdU (mitosis and meiosis S phase) and TRA98 (germ cell marker). (B) Representative photomicrographs of SYCP3 (green), BrdU (red) and TRA98 (blue) immunostaining. Scale bar 5 µm. Pre PL = pre-leptotene; post PL= post-leptotene. Error bars show mean ± s.e.m. n=5 mice from independent exposures; * p ≤ 0.05; ** p ≤ 0.01 (Mann-Whitney’s test). (C) Graphs showing the percentage of preleptotene (BrdU and SYCP3 positive) and meiotic (BrdU negative, SYCP3 positive) oocytes. Error bars show mean ± s.e.m. n=5 mice from independent exposures; * *p* < 0.05; ** *p* < 0.01 (Mann-Whitney’s test).

This defect in meiosis progression could be the consequence of a delay in meiosis initiation. Immunostainings for BrdU, SYCP3 and a germ cells marker (TRA98), were used to identify three germinal populations 14.5 dpc ovaries: oogonia, the female PGCs (SYCP3-negative cells), premeiotic cells in S-phase also called pre-leptonema (BrdU-positive/SYCP3-positive) and oocytes (BrdU-negative/SYCP3-positive; Figure 3B-C). In untreated 14.5 dpc ovaries almost all germ cells had initiated meiosis (over 96% are SYCP3+) and very few oogonia and preleptotene cells were still present. In bisphenols-treated mice, we observed a significant increase in mitotic PGCs (Supplementary Figure 1A) and pre-leptonema while oocyte number was reduced (Figure 3C).

In addition, we observed an increasing trend for STRA8 positive germ cells in 14.5 dpc female gonads (20% ± 4 in ETOH group *VS* 36% ± 1 *p*=0.08 and 39% ± 6, *p*=0.07 in BADGE and BPAF groups respectively, Supplementary Figure 1B). The presence of STRA8 correlated with the initiation of the meiotic program and declines rapidly just after the initiation in prophase I. The observed increase confirmed the bisphenols-induced delay of meiotic initiation. A defect of crossover distribution could be the consequence of a delay and/or an alteration of meiosis initiation. To confirm this hypothesis, we performed exposure to bisphenols until meiosis initiation (short term exposure).

Fetuses were exposed from 10.5 to 14.5 dpc (Supplementary Figure 2, upper panel) and the MLH1 foci was quantified at 18.5 dpc. As observed for bisphenols exposures during meiosis progression (long-term exposure), the number of MLH1 foci was significantly increased after bisphenols exposure (Supplementary Figure 2). This suggests that changes in crossovers distribution were probably due to alterations occurring before or during mitosis to meiosis transition in germ cells (GCs).

### BADGE or BPAF fetal exposures alter gene expression and mRNA splicing in germ cells

In order to understand the bases of bisphenols-induced alterations in PGCs during acquisition of meiotic competence and initiation of the meiotic program, we performed transcripomic analyses on 11.5 dpc (during acquisition of the meiotic competence) and 13.5 dpc (during meiosis initiation) germ cells. We sorted PGCs by Magnetic Activated Cell Sorting (MACS) using the cell surface protein stage-specific embryonic antigen 1 (SSEA-1). Gene expression analysis was conducted using murine Affymetrix GeneChip Gene 2.0 TS (11.5 dpc) and Clariom™ D (13.5 dpc) microarrays. At 11.5 dpc, we identified 1817 and 733 differentially expressed genes (DEGs) in BADGE and BPAF treated germ cells respectively (compared to vehicle treated PGCs with a |Log2 Fold-Change| ≥ 0.5 and *p* < 0.05) (Figure 4). DEGs were mostly downregulated (56 to 65 % of DEGs; Figure 4A). At 13.5 dpc when the program of meiosis is initiated, we identified 886 and 1376 DEGs in BADGE and BPAF treated germ cells respectively (Figure 4A). Contrary to what it is observed at 11.5 dpc, two third of the DEGs were upregulated in 13.5 dpc bisphenols-treated germ cells (Figure 4A). Using EnrichGO function from Clusterprofiler package to identify enrichment of gene ontologies, we observed that DEGs at 11.5 dpc as well as 13.5 dpc were mostly related to meiosis, stem cell differentiation and regulation of stem cell signaling pathway such as Wnt pathway (Figure 4B and Supplementary Figure 3). Meiosis-associated DEGs such as *Stag3, Sycp1*&*3, Hormad1*&*2, Spo11, Spata22, Meiob, Dmc1* or *Brca2* were strongly enriched in genes associated to synapsis and DNA recombination processes (Figure 4A-C, Supplementary Table 1). Surprisingly, genes linked to meiosis and stem cell differentiation showed opposite transcriptional response at 11.5 dpc and 13.5 dpc (Supplementary Figure 3 and Figure 4C). Stemness genes tend to be up-regulated at 11.5 dpc and mostly down-regulated at 13.5 dpc and meiotic genes were preferentially down-regulated at 11.5 dpc and up-regulated at 13.5 dpc (Figure 4C).

**Figure 4:**
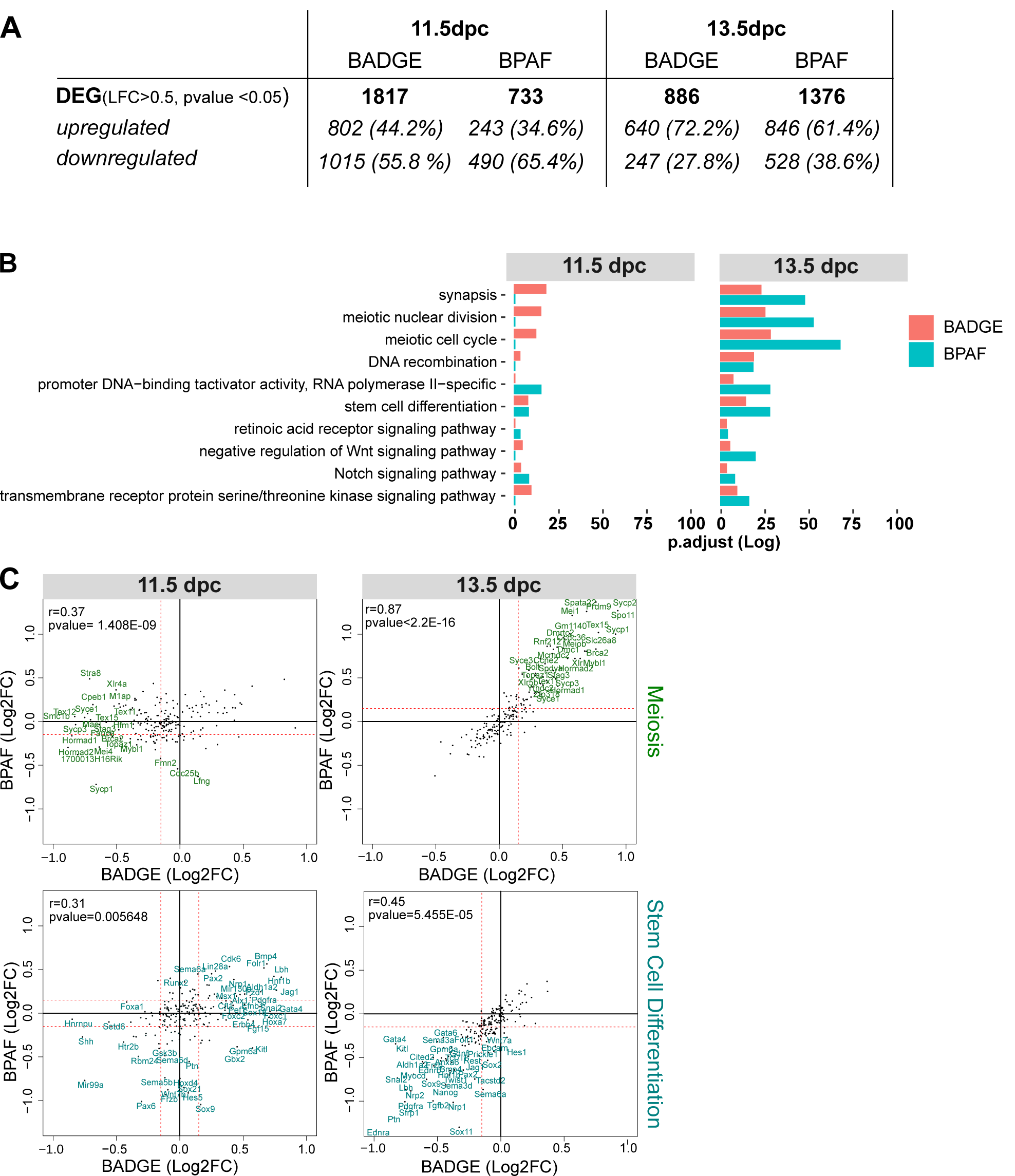
BADGE or BPAF fetal exposures induce transcriptional alteration before meiosis initiation. SSEA1 positive germ cells from 11.5 or 13.5 dpc ovaries treated by vehicle (ETOH) or bisphenols (BADGE and BPAF) were used for transcriptomic analyses. Differentially expressed genes (DEGs) between bisphenols and control condition (BPAF vs ETOH or BADGE vs ETOH) were filtered according to the log fold change (*abs*[Log2FC] ≥ 0.5) and the significativity (*p* < 0.05). Three pools of cells each of them from 10-15 fetuses were analyzed. (A) The table represents the number (and percentage) of DEGs after BADGE or BPAF exposure in 11.5 or 13.5 dpc germ cells. (B) Gene Ontology enrichment associated to meiosis and stem cell differentiation among DEGs. (C) Correlation analysis between BADGE and BPAF condition of the differential expression (Log2FC) of genes associated to meiosis or stem cell differentiation. *r*= Pearson’s correlation coefficient (Pearson’s test).

Interestingly, we observed a significant correlation between BPAF and BADGE of the differential expression of genes associated to meiosis after bisphenols exposure (r=0.37, *pvalue <* 0.0001, 11.5dpc and r=0.87, *pvalue* < 0.0001, 13.5 dpc) and stem cell (r=0.32, *p<*0.0001, 11.5 dpc and r=0.45, *p* < 0.0001, 13.5 dpc) at 11.5 dpc and 13.5 dpc (Figure 4C). These positive correlations demonstrate a common transcriptional signature between BPAF and BADGE exposure and suggest similar mechanisms of action for both bisphenols. Interestingly, as observed in the chromosomes 3, 4, 7, 10, 14, and 17, genome mapping of DEGs (up- and down-regulated) showed a concentration of genes on specific genome locations that were differentially expressed after BADGE and BPAF exposure at 11.5 dpc as well as 13.5 dpc (Supplementary Figure 5). A regulated alternative splicing program has been reported during meiosis initiation in male and in female germ cells (Naro et al., 2017; J. Wang et al., 2019). The exact role of this program is unclear but is though to be required for stabilization of meiotic transcripts during prophase I of meiosis (Liu et al., 2017; J. Wang et al., 2019). As observed in adult testis, modulation of this program results in spermatogenesis defects, highlighting the essential role of the RNA splicing during meiosis (Hannigan et al., 2017; Liu et al., 2017). For this reason, we studied the impact of bisphenols exposure on RNA splicing events during meiosis initiation. We took advantage of the Clariom™ D that allows the detection of most RNA splicing events by covering independently exons and exons junctions of all transcripts. After bisphenols exposure, 3543 (BADGE) and 2678 (BPAF) coding or non-coding genes showed one or more alternative RNA splicing events compared to the control condition (Figure 5A-C). Differentially spliced genes were strongly associated with chromosome segregation, meiotic cell cycle and DNA repair and recombination (Figure 5A). Half of them were identical in BADGE and BPAF -treated germ cells including genes such as *Smc4, Sun1, Mcmdc2, Dazl, Rad51, Mlh1, Dmc1, Sycp2* (Table 1 and Supplementary Table 2). Different modes of alternative splicing including skipped exons, alternative 5′ splice sites, alternative 3′ splice sites, and retained intron are described in metazoans. TAC software classifies some of these events in function of the coverage of the probes sets of the array. Bisphenols-treated germ cells showed an enrichment of transcripts with skipped exon features in all genes including meiotic specific genes (Figure 5B, Table 2).

**Table 1:**
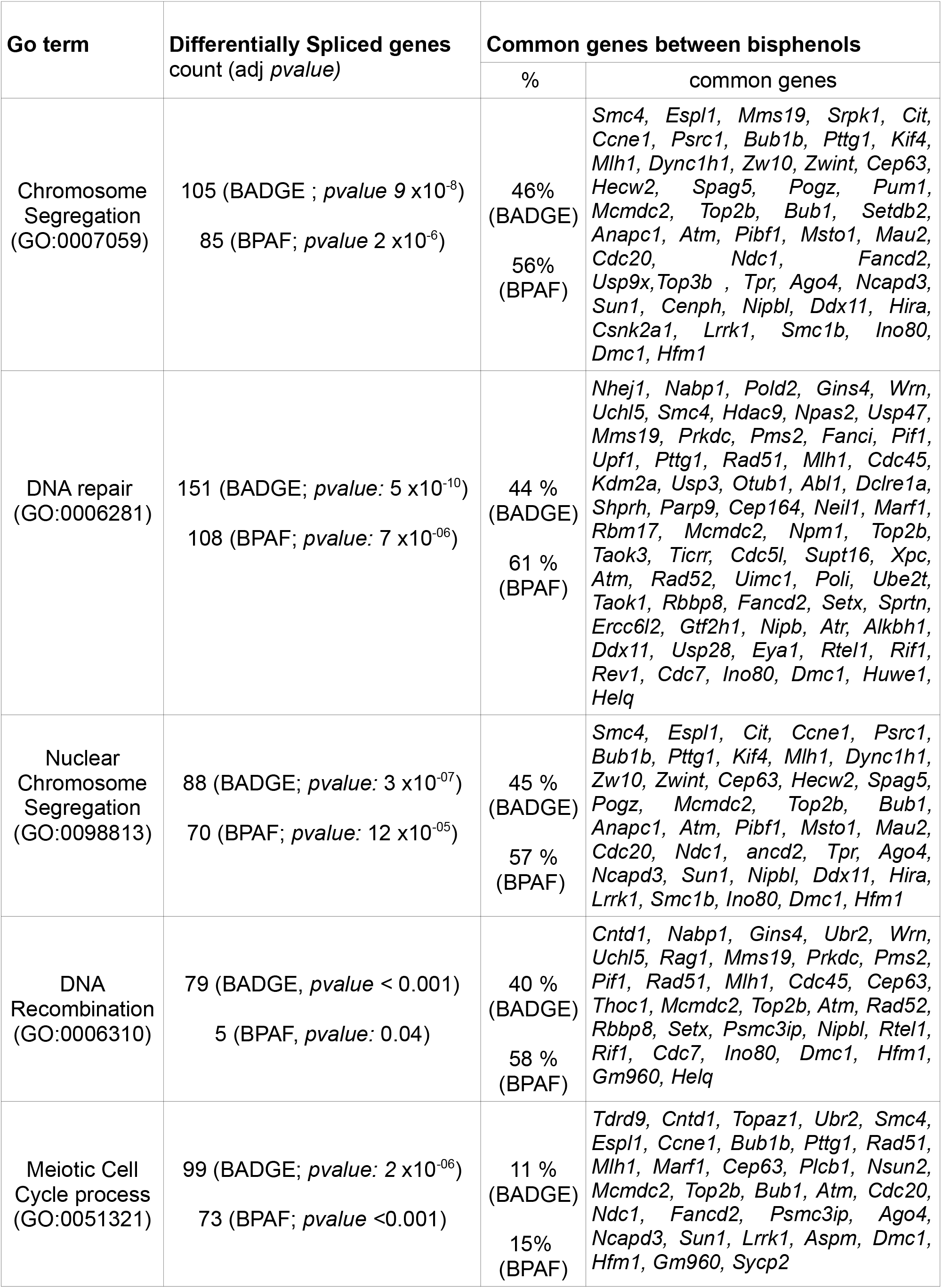
List of common genes between BADGE and BPAF conditions showing an alternative spllcing relative to control germ cell.

**Table 2:**
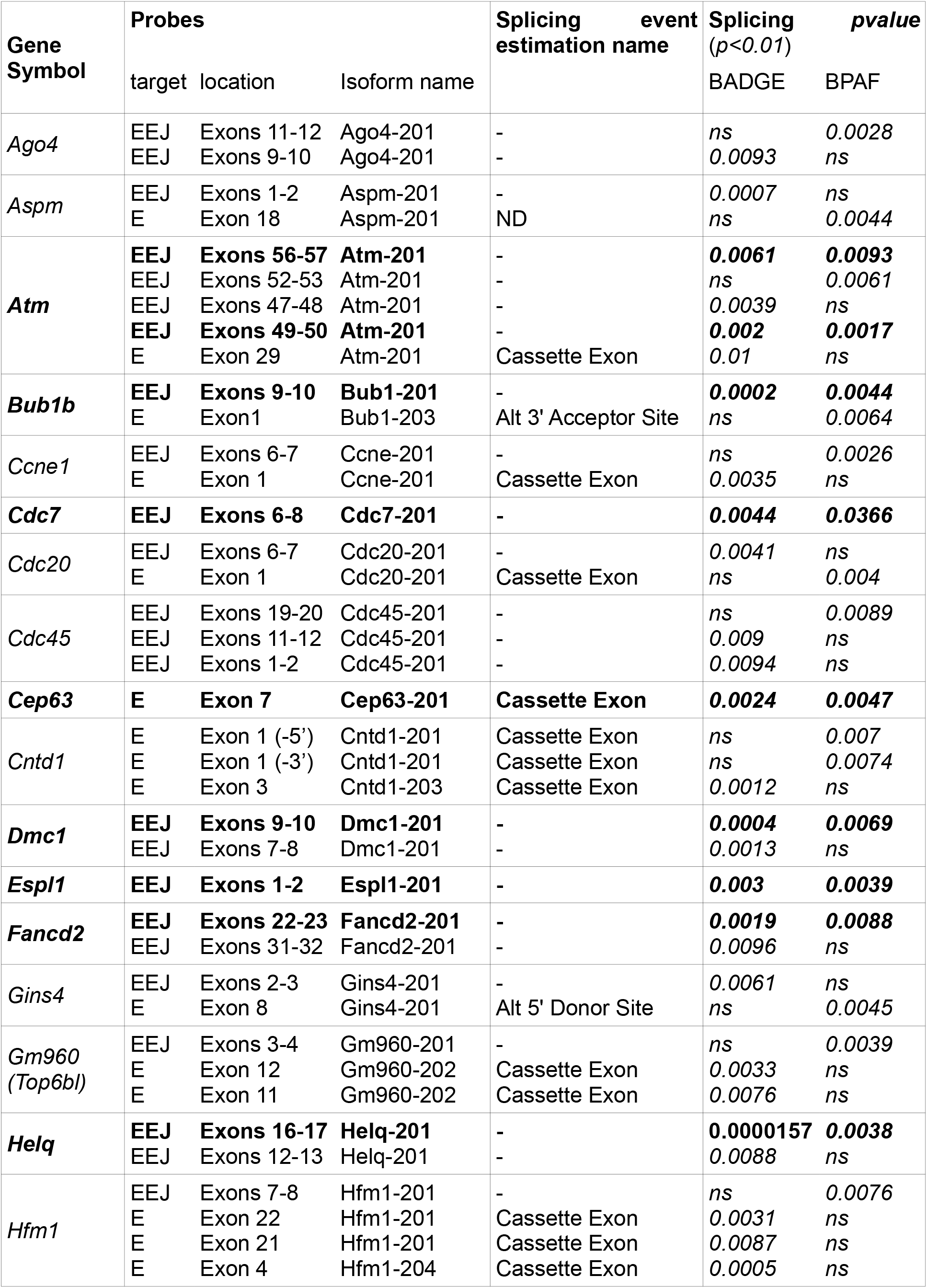
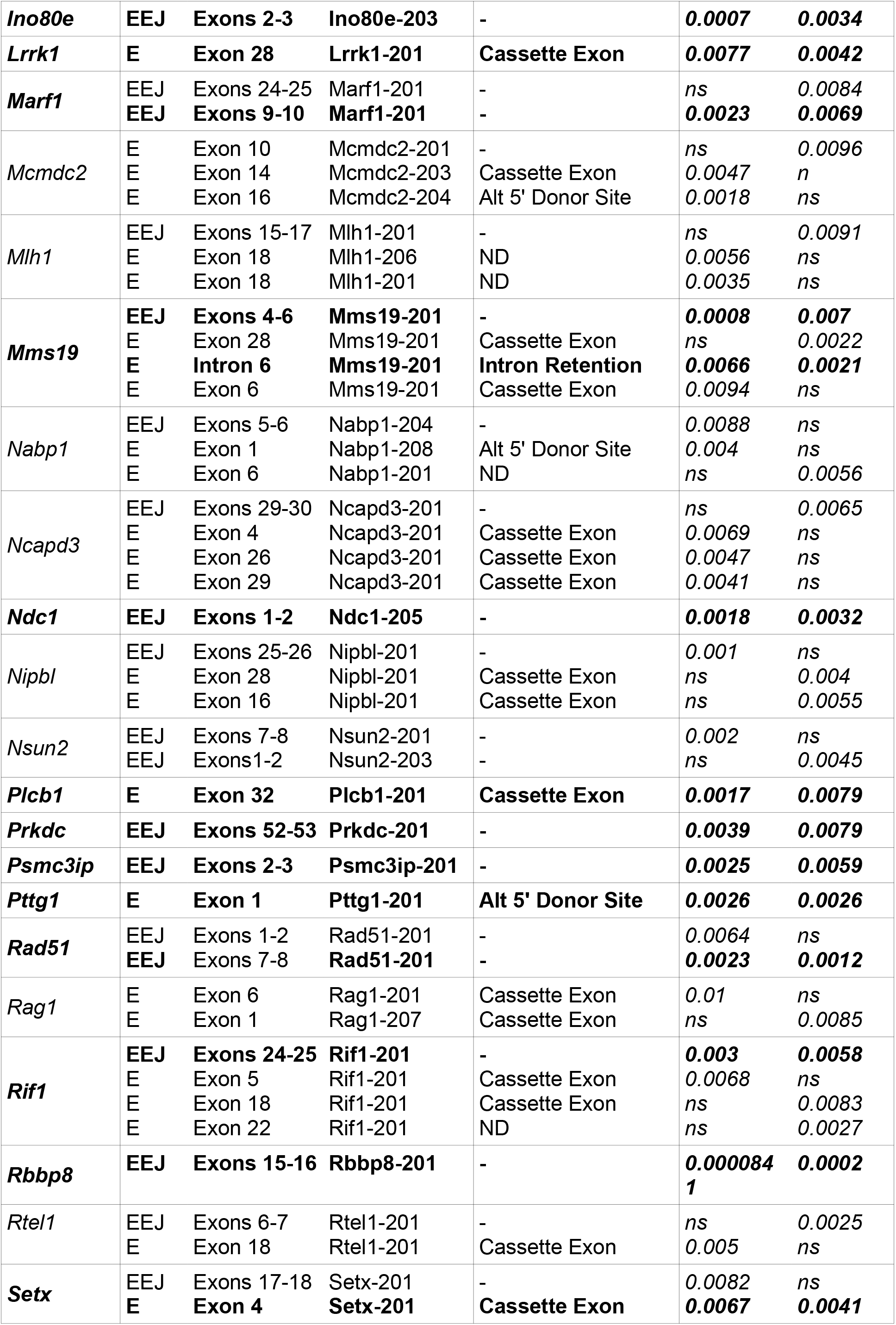
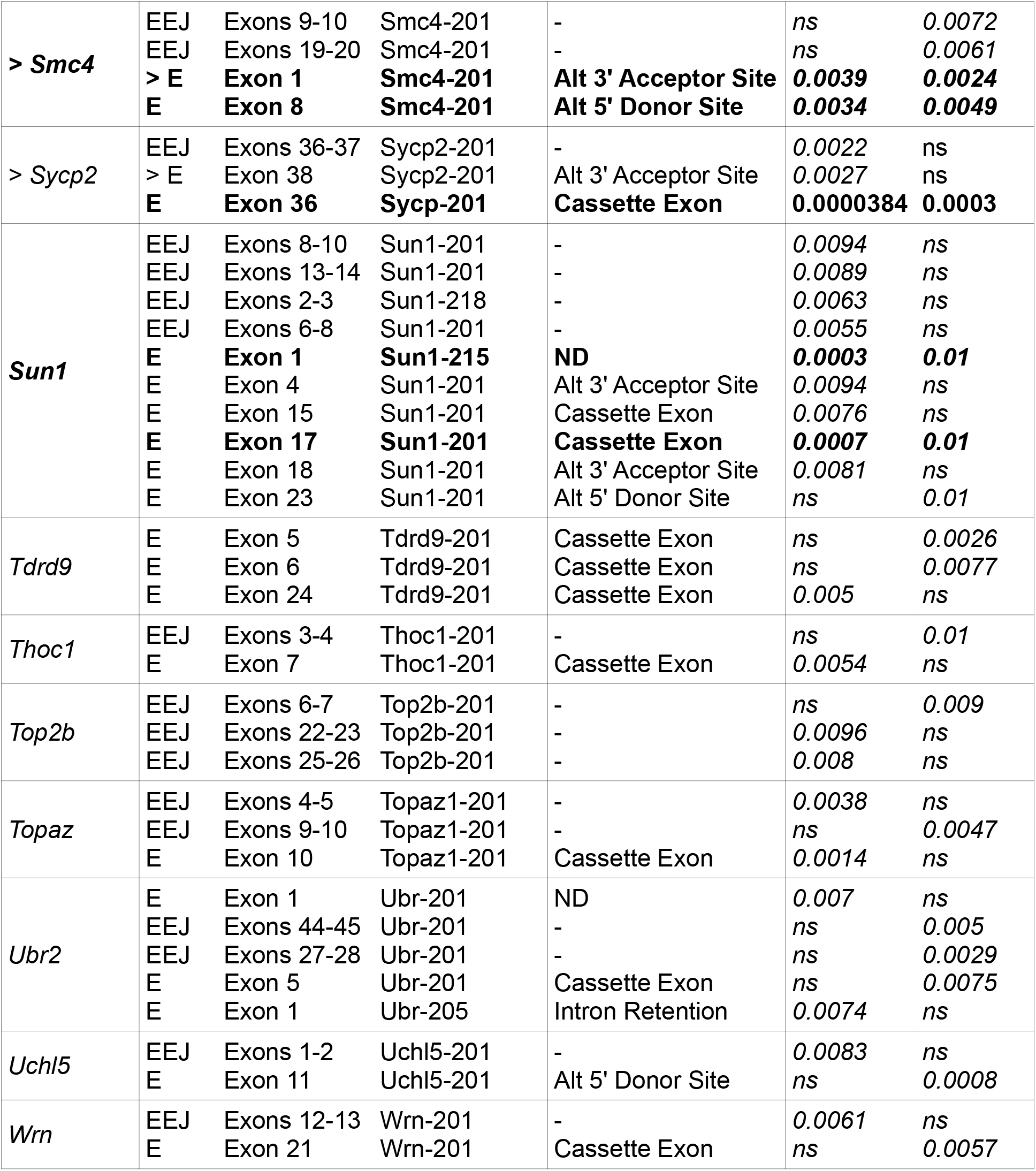
Splicing event estimation by TAC software in bisphenols treated germ cells. EEJ: one exon-exon junction was covered by probes, E: one exon was covered by probes. Only PSRs have assigned « Event Estimation Name ». Embedded text: common significative splicing events between BADGE and BPAF.

**Figure 5:**
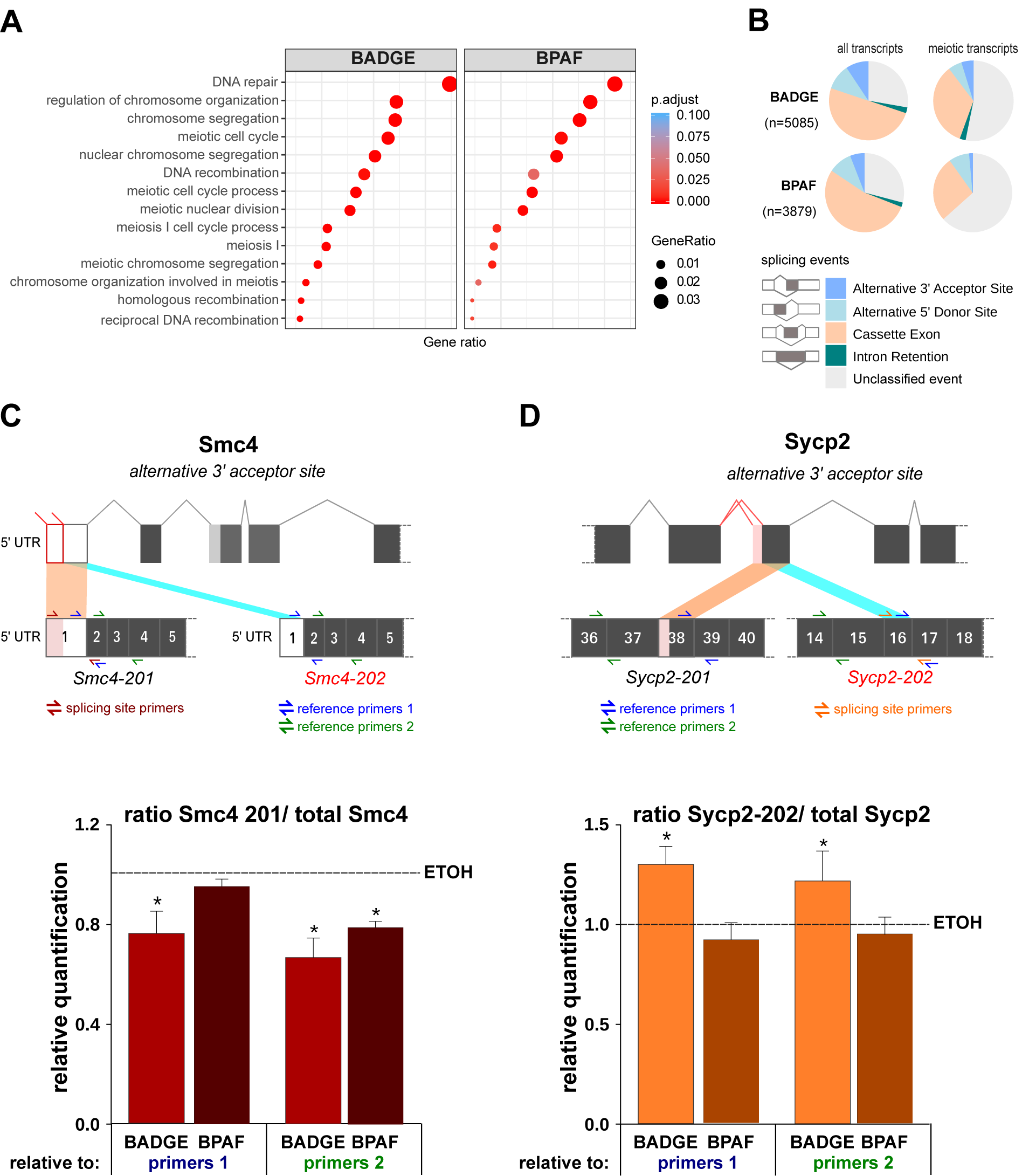
BADGE or BPAF fetal exposures induce RNA splicing alterations in premeiotic germ cells. Splicing events were determined in 13.5 dpc germ cells using the high resolutive microarray. Using TAC software, differentially spliced genes (BPAF vs ETOH and BADGE vs ETOH) were filtered according to the exon splicing index (*abs*[exon splicing index]≥1) and the exon *pvalue* (*p*<0.05).n=3 pools of cells from 10-15 fetuses each. (A) dot plot showing enrichment on ontology associated to meiosis of differentially spliced genes. (B) distribution of splicing events in all genes and specifically meiotic genes. Modes of alternative splicing including skipped exons, alternative 5′ splice sites, alternative 3′ splice sites, and retained intron. Unclassified events correspond to events that cannot be included in any of the standard categories. **(C) and (D)** Quantification of the relative incidence of *Smc4* **(C) and** *Sycp2* (D) splice variants (Ensembl variants:) in bisphenols-treated germ cells. n=3 independent exposures. Upper panel: Representation of splicing sites leading to different splice variants (*Smc4-201, Smc4-202, Sycp2-201* and *Sycp2-202*, Ensembl name). Lower pannel: relative quantification of expression of *Smc4-201 or Sycp2-202* to total *Smc4* transcripts and *Sycp2* transcripts respectively in bisphenols treated germ cells in comparison to ETOH-treated germ cells. * *p* <0.05 (Mann-Whitney’s test).

Some of the differentially spliced meiotic genes retrieved in both bisphenols exhibit the same splicing events such as *Smc4*, a condensin protein (Table 2 in red, Figure 5C). The presence of an alternative 3’ acceptor sites in the first exon allows the production of different forms of *Smc4* transcripts containing or not a truncated exon 1 as observed in the *Smc4-202* and *Smc4-201* variants respectively. Using specific primers of the truncated/spliced region of the first exon and of the core/unspliced region of the transcript, we quantified by QPCR the relative abundance of the mRNAs containing a long first exon. In BADGE as in BPAF -treated germ cells, we observed a significant decrease in the abundance of the long form with a corresponding increase of the truncated form of this first exon (Figure 5C). While we retrieved the same affected genes in BADGE and BPAF treated germ cells, the precise differential splicing events differs between conditions as observe for meiotic gene set (Table 2 in black). In BADGE -treated germ cells, we identified a differentially specific event occurring in the exon 38 of *Sycp2*, a component of the synaptonemal complex, due to alternative 3’ acceptor sites (Table 2). These sites lead to the production of two types of variants with a complete or a truncated exon as observed in *Sycp2-201* and *Sycp2-202* transcripts. QPCR quantification revealed a significant increase in the relative abundance of the truncated form of the exon 38 only after BADGE exposure (Figure 5D).

Genes differentially spliced in BPAF or BADGE conditions were not preferentially differentially expressed compared to others genes (Supplementary Figure 4). As observed with non-spliced genes, less than 8 % of spliced genes were also differentially expressed in BPAF or BADGE conditions. Interestingly, we observed closer transcription behaviors after BPAF or BADGE exposure of differentially spliced genes in comparison to non-differentially spliced genes randomly chosen (Supplementary Figure 4). This observation suggests a common alteration impacting the transcriptional response and RNA splicing after BADGE and BPAF exposure.

### BADGE or BPAF fetal exposures increase oxidative DNA damages impairing meiosis initiation

As BPA is a well-known oxidative stress generator in germ cells and other cell types we speculated that the formation of oxidative DNA damage such as 8-OdG might explain common transcriptional responses and RNA splicing after BADGE or BPAF exposure. Indeed, it is well known that the formation of oxidative DNA damages such as 8-OdG alters gene expression (Ba et Boldogh 2018). Using Oxidip-seq data obtained by Amente et al (Amente et al. 2019) on mouse embryonic fibroblasts (MEF), we observed that DEGs genes could overlap with DNA regions susceptible to be oxidized (Supplementary Figure 5).

Therefore, we analyzed the induction of oxidative DNA damages in response to bisphenols in 12.5 dpc proliferative oogonia by detection of 8-OdG (Figure 6A-B). As a positive control, we added the pro-oxidant potassium bromate (KBrO3) in drinking water from 10.5 dpc. As observed after KBrO3 exposure, bisphenols-exposed PGCs showed a significant increase in 8-OdG when compared to the vehicle-treated germ cells (Figure 6B).

**Figure 6:**
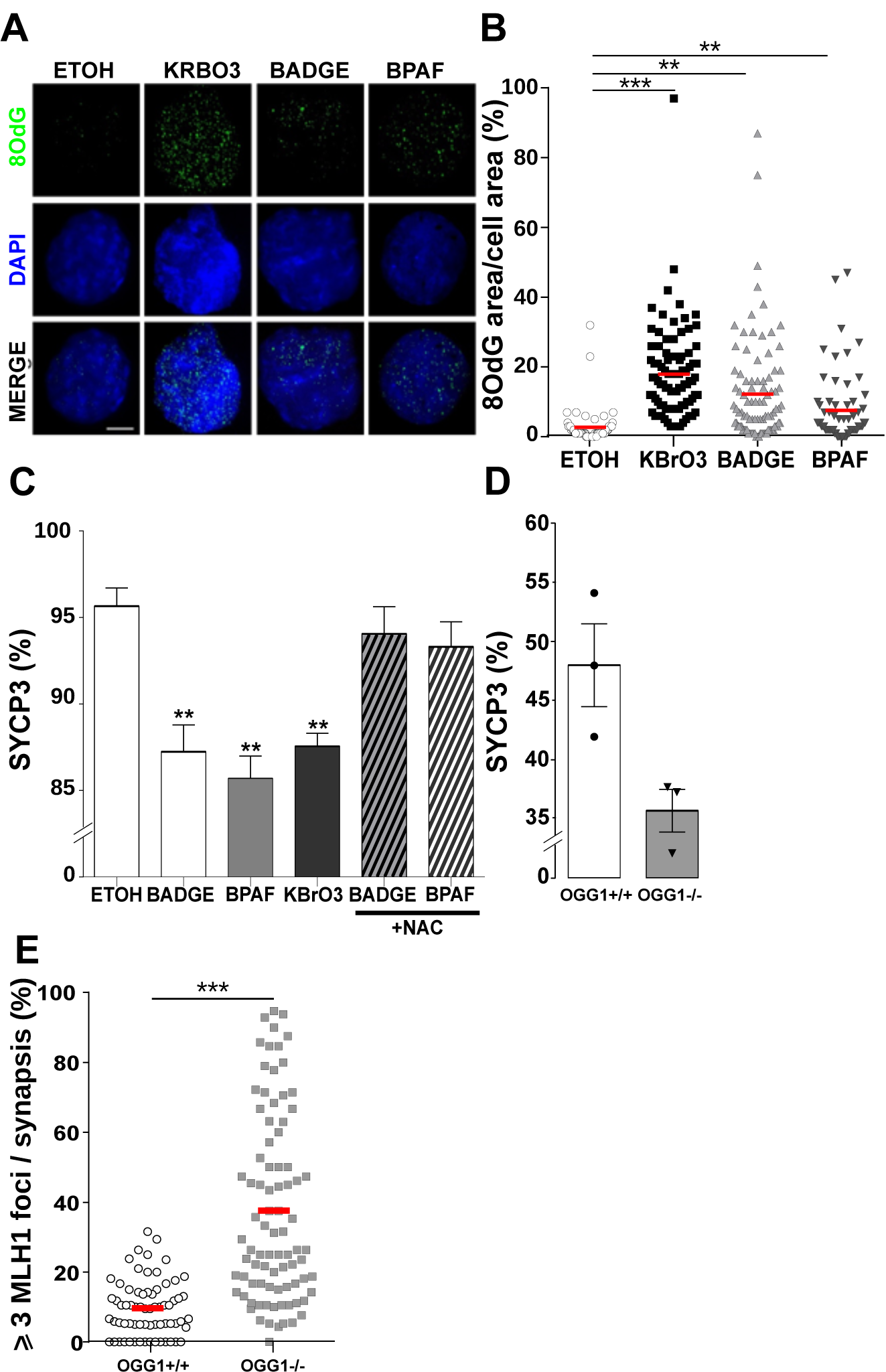
Oxidative DNA damages induced by bisphenols impact meiosis initiation and meiotic recombination. (A) 8-oxoguanine (8OdG, green) immunostaining in 12.5 dpc germ cells (TRA98 positive cells) from ovaries exposed to vehicle (ETOH), bisphenols (BADGE, BPAF) or the pro-oxidant KbrO3 (0.15 mM). Scale bar, 10µm. (B) Quantification of the total area of 8OdG normalized by the nucleus area. n= 60 (ETOH), n=90 (KrBO3), n=94 (BADGE) and 64 (BPAF) oogonia from 3-4 independant exposures. (C) Percentage of meiotic oocytes in germ cells identified by SYCP3 and TRA98 immunostaining in 14.5 dpc ovaries exposed to vehicle (ETOH), KrBO3 (0.15 mM) and bisphenols (BADGE, BPAF) supplemented with or without NAC (4 mM). Error bars show mean±s.e.m. n=3-5 mice from independant exposures; ** p<0.01; *** p<0.001; **** p<0.0001 (Mann-Whitney’s test). (D) Percentage of meiotic oocytes in 14.5 dpc germ cells from OGG1 deficient (-/-) and wild-type (+/+) mice. n=3-4 mice. *p= 0*.*0832* determined with a Mann-Whitney’s test. (E) Percentage of synaptonemal complexes with 3 or more MLH1 foci in pachytene cells identified on the basis of the SYCP3 in OGG1 deficient (-/-) and wild-type (+/+) 18.5 dpc fetuses. n= 64 (OGG^+/+^), n=80 (OGG^-/-^) from 3 fetuses. *** p<0,001 (Mann-Whitney’s test).

To explore the relationship between meiosis initiation delay and oxidative stress, we quantified meiosis initiation in KBrO3 treated ovaries at 14.5 dpc. KBrO3 exposure significantly decreased the proportion of meiotic oocytes (SYCP3 positive cells; Figure 6C).

On the contrary, when we treated pregnant mice with bisphenols supplemented with the antioxidant N-acetylcysteine (NAC) to pregnant mice, we restored the percentage of oocytes to that of vehicle treated (ETOH) 14.5 dpc ovaries. To further asses the impact of oxidative lesions, we used mice deficient for OGG1, a DNA glycosylase involved in the removal of oxidative DNA damage through the Base Excision Repair (BER). *Ogg1*^-/-^ mice showed increased levels of 8-OdG in proliferative 12.5 dpc oogonia in comparison to *Ogg1*^+/+^ mice (Supplementary Figure 6). The lack of OGG1 also impaired meiosis initiation as we observed a decreasing trend of oocyte number at 14.5 dpc (48.0% ± 3.5, *Ogg1*^+/+^ vs 35.7% ± 1.8, *Ogg1*^-/-^, p=0.0832; Figure 6D). To link oxidative DNA damage and recombination events, we quantified the number of MLH1 foci at 18.5 dpc in OGG1 deficient mice. Interestingly, as observed for bisphenols treated mice, the number of MLH1 foci was significantly increased in the mutant when compared to wild type mice (Figure 6E). Taken together these data reveal that bisphenols induce oxidative DNA damages which in turn could delay fetal meiosis and induce recombination defects.

### BADGE or BPAF fetal exposures alter DNA demethylation in PGCs

BPA has been reported to alter DNA methylation in various cell types and removal of DNA methylation is a mandatory event in proliferative PGC to allow later the proper expression of the meiotic program(Anderson et al., 2012; Chao et al., 2012; Hargan-Calvopina et al., 2016b; Yamaguchi et al., 2012). For this reason, we quantified the proportion of methylated/pluripotent PGC compared to unmethylated/differentiating PGC by immunodetection in 12.5 dpc bisphenols exposed and unexposed ovaries of the 5-hydroxymethylCytosine (5hmC), the first intermediate form during active demethylation (Figure 7A). After bisphenols exposure, we observed an enrichment of methylated germ cells with a global staining (homogeneous staining) of 5hmC in the nucleus in TRA98 cells (Figure 7B). This result suggests that bisphenols exposure alters or delays the DNA demethylation in PGC increasing the proportion of methylated germ cells compared to unmethylated ones.

**Figure 7:**
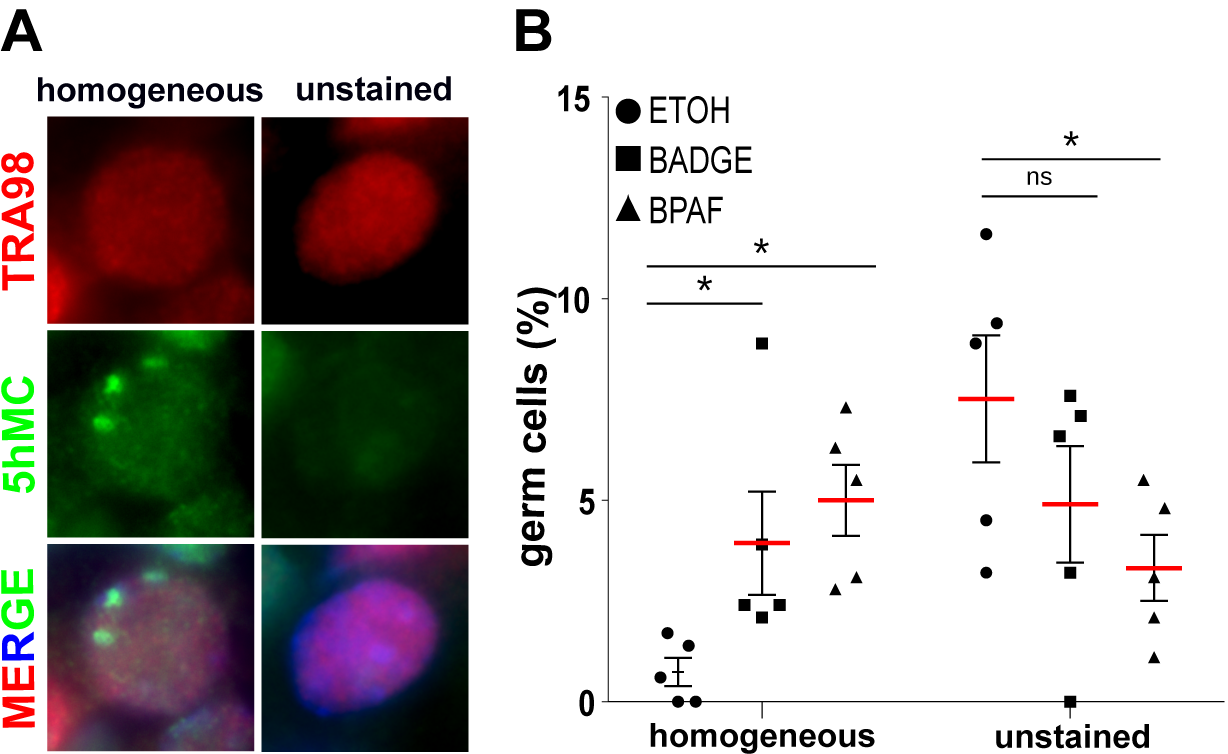
BADGE or BPAF fetal exposures alters DNA methylation in proliferative PGC. Immunodetection of 5hmC (green) in 12.5 dpc TRA98 positive PGC (red) counterstained with DAPI (blue). (A) Representative photomicrograph of methylated (homogeneous nuclear staining) and unmethylated (unstained) TRA98 PGC. (B) percentage of methylated and unmethylated PGC. n= 5 from 3 different exposures. * p<0.05 (Mann-Whitney’s test).

## Discussion

Woman’s reproductive pathologies include the ovarian dysgenesis syndrome that is defined as a precocious alteration of ovarian function and mostly associated with premature ovarian insufficiency, anovulation and/or pregnancy loss in the adulthood (Buck Louis et al., 2011). Oocyte aneuploidy is strongly associated with these reproductive failures and growing evidences suggest that fetal exposure to endocrine disruptors such as Bisphenol A (BPA) can cause female reproductive defect (Johansson et al., 2017). In Vertebrates and Invertebrates species, exposure to BPA induces transcription defect of meiotic genes, alters frequency of MLH1 foci and chiasmata leading to chromosome missegregation during meiotic resumption (Allard & Colaiácovo, 2010a; Brieno-Enriquez et al., 2012; Brieño-Enríquez et al., 2011; P. A. Hunt et al., 2012; Patricia A. Hunt et al., 2003; Lawson et al., 2011; Susiarjo et al., 2007). Because BPA has been restricted and regulated in many countries, BPA alternatives have been introduced into industrial products increasing the exposure to other bisphenols such as BPAF and BADGE. Exposure to BADGE and BPAF can be comparable to BPA exposure. A recent study has reported comparable median concentrations of BPA and BADGE hydrated derivatives in human plasma collected from New York City individuals (7.15 ng/ml for BADGE·2H20 and 2.26 ng/ml for BADGE·H20 versus 1.77 ng/ml for BPA) likely due to similar exposure source to BPA (ie. from epoxy coatings into canned food) (L. Wang et al., 2015). In China, BPAF mean concentration in human plasma is 0.073 ng/mL versus 0.4 ng/ml for BPA in the same samples from 81 individuals in general population (Jin et al., 2018). For this reason, we investigated the consequences of BPAF and BADGE exposure on female germ cell differentiation. Although BPAF concentration detected in human samples is lower than the one of BADGE or BPA, we chose to expose mice by drinking water to BADGE or BPAF at the same concentrations (*ie* 10µM). This allows to compare BPAF and BADGE and to compare our results to previous studies, including ours, that used BPA at environmentally doses (Eladak et al., 2018). In the present study, we exposed pregnant mice to bisphenols from 10.5 to 18.5 dpc. The concentration of total BPAF detected in the plasma of the pregnant mice was of the a same order of magnitude (5.96 ± 0.78 ng/ml) than the one we previously observed for BPA (Eladak et al., 2018).

As observed for BPA, fetal exposure to BPAF or BADGE induces subtle oocyte defects during fetal life leading to aneuploidy at adulthood. We observed changes in expression levels of meiotic genes during meiosis initiation, and a modification of the frequency of MLH1 foci during pachynema and of chiasmata in diplonema leading to oocyte aneuploidy after meiotic division. Thus, we demonstrate that BPAF or BADGE seem as much harmful as BPA with regard to their deleterious effect on oogenesis.

Until now the modality of the action of BPA on meiosis remains unclear. Numerous evidences highlight a temporal window of oocyte vulnerability during fetal life, specifically before and during meiotic prophase I (Brieno-Enriquez et al., 2012; P. A. Hunt et al., 2012; Lawson et al., 2011; Susiarjo et al., 2007). Because consequences are mostly detectable in later steps it was difficult to determine the precise cellular target (*ie*. PGC or oocyte) of BPA. This questionned whether the bisphenol-induced meiotic failure originated from alterations that occurred during the differentiation of the PGC or during meiotic prophase I. In this study, we showed that exposure to bisphenols before meiotic prophase I affected crossover frequency. Therefore, this observation demonstrates that alterations induced by BPAF and BADGE in PGC alters the establishment of the meiotic program.

Proper establishment and orchestration of the meiotic program are essential to ensure a correct regulation of crossover distribution [57]. The establishment of prophase I requires two events: acquisition of meiotic competence and initiation of the meiotic program (Soh et al., 2015). Meiotic competence is acquired after PGCs colonization. Migratory PGCs have a genomic program associated with stemness and express pluripotent genes. In the gonad, at 10.5 dpc, PGCs switch off the pluripotency program and activate genes involved in gametogenesis. DAZL, a germ cell-specific RNA binding protein, binds germ cell specific mRNA and promotes the stabilization of meiotic mRNA allowing the initiation of the meiotic program (Atala, 2012; Hu et al., 2015; Kato et al., 2016; Nicholls et al., 2019). Our transcriptomic analyses revealed that bisphenol exposure alters the acquisition of the gametogenic competence and the initiation of the meiotic program. First, bisphenol exposure seems to induce a failure or a delay on the switch from pluripotency to gametogenesis/meiosis competence. This is illustrated by a global down-regulation of meiosis associated genes such as *Sycp1&3* and *Hormad2* and an up-regulation of pluripotency associated genes such as *Msx1* and *Gata4*. Second, bisphenols exposure also impacts the initiation of the meiotic program at 13.5 dpc as illustrated by the up-regulation of meiotic genes. These observations are consistent with previous studies on the effect of BPA on human and murine meiosis initiation (Houmard et al., 2009; Lawson et al., 2011). However, it is unclear whether the up-regulation of meiotic genes induced by bisphenols is the consequence of the PGC differentiation delay observed earlier, or a real induction of gene expression and/or RNA stabilization. Indeed, numerous meiotic genes such as *Stra8* or *Rec8* are drastically down-regulated just after meiosis onset. Therefore, a delay of meiosis initiation would lead us to observe a higher level of expression of some meiotic transcripts (Soh et al., 2015). Moreover, bisphenol exposure also induced a global alteration of splicing events during this step. Recent studies have highlighted the role of mRNA splicing on the mitosis to meiosis transition in male and female germ cells and lack of splicing regulators in male germ cells leads to meiotic defects (Liu et al., 2017; Naro et al., 2017; Schmid et al., 2013; J. Wang et al., 2019). In consequences, minor modifications of the splicing of meiotic genes after bisphenols exposure could impact meiosis. As observed for BPA exposure, changes in level of gene expression were subtle but, interestingly, we observed common transcriptional signature between BPAF and BADGE suggesting common bisphenol molecular targets.

Transcriptional modifications observed after bisphenol exposure impacted the transition from mitosis to meiosis and were certainly the origin of the observed delay of meiosis initiation and progression, the abnormal distribution of MLH1 foci but also the cause of folliculogenesis alterations (ie.MOF, follicle number). Indeed, the initiation of follicle assembly requires completion of meiotic prophase and any delay in meiotic progression could interfere with this process (Chao et al., 2012; A. Paredes et al., 2005; H.-Q. Zhang et al., 2012). Thus all these events may be related forming a cascade resulting from the early alteration of the establishment of the meiotic program in PGC.

Interestingly, common cellular (*ie* MLH1 and chiasmata distribution and MOF) and transcriptional meiotic signatures observed after bisphenols exposure and other pollutants such as phthalates in multiple organisms suggest a common mechanism of action of these molecules (Allard & Colaiácovo, 2010b; Cuenca et al., 2020; Gely-Pernot et al., 2017; Parodi et al., 2015; Shin et al., 2019; Susiarjo et al., 2007; Tu et al., 2019). In this study, we propose oxidative stress as an new player involved in bisphenol response that could impair the PGC differentiation. One of the major targets of reactive oxygen species is the DNA, and 8OdG is the most prominent lesion in the genome (Ba & Boldogh, 2018). We showed that bisphenol exposure quickly induces the formation of this oxidative DNA lesion in proliferative germ cells. Interestingly, analyzing the genomic landscape of oxidative DNA damage revealed a specific susceptibility to oxidation in genomic regions harboring genes differentially expressed after bisphenol exposure. It is well known that oxidative DNA damages can modulate gene expression directly or undirectly. First, oxidative DNA damage induce antioxidant and inflammatory transcriptional responses and recruiting DNA repair proteins (Hörandl & Hadacek, 2013; Pan et al., 2016). Second, oxidized guanine biases the recognition of methylated CpG dinucleotides and alters the dynamic of DNA demethylation (Gruber et al., 2018; Pan et al., 2016). Interestingly, acquisition of the meiotic competence is completely dependent on an intensive germinal DNA demethylation (Hargan-Calvopina et al., 2016b; Yamaguchi et al., 2012). In this study, we observed that bisphenols exposure interferes with DNA demethylation and could be the consequence of the presence of oxidative DNA damages. The presence of 8OdG has also post-transcriptional consequences and could alter splicing events. Pre-mRNA splicing requires the identification of specific 5’ donor and 3’ acceptor sites. The 5’ and 3’ splicing sites mostly begin with dinucleotide GT(U) and end with dinucleotide AG that allows the major 5’_3’ combination GT-AG (Calvello et al., 2013). Incorporation of the wrong base within these splicing signals would lead to an alteration of the fidelity of mRNA splicing. In mammalian cells, unrepaired DNA lesions such as 8OdG on the transcribed strand is a source of transcriptional mutagenesis. Unrepaired 8OdG leads to a single-nucleotide substitution in the 3’ or 5’ splice site subsequently resulting mostly in exon skipping and altering the protein product as it has been observed in the context of Ogg1 deficiency (J. A. Paredes et al., 2017). For these reasons, we speculated that induction of oxidative stress by bisphenols could be responsible of observed transcriptional alteration as well as RNA splicing modification during meiosis initiation leading to prophase I defects. This is confirmed after specific induction of oxidative DNA damages in proliferative germ cell. Indeed, in KrBO3-treated ovaries and the timing of meiosis initiation is restored in presence of the antioxidant NAC. Moreover, in Ogg1 deficient mice, we also observed a delay of meiosis initiation and modification of MLH1 distribution in pachynema.

This study highlights the central role of oxidative DNA damage in the meiotic response after bisphenol exposure. Numerous past studies proposed a role for estrogen signaling in the meiotic responses (*ie* delay of meiotic progression, alteration of crossover distribution and aneuploidy) due to the effect of molecules with a xenoestrogenic potential (Cuenca et al., 2020; Susiarjo et al., 2007; Tu et al., 2019). Our hypothesis is in agreement with an involvement of estrogen signaling during this process as DNA binding by the estrogen receptor drives the local production of oxidative DNA damages via LSD1, a Lysine-specific histone demethylase 1, activity in promoter and enhancer regions (Perillo et al., 2008).

Current knowledge on the potential toxicological effects of BPA analogs is limited. We provide here proofs that endocrine disruptors such as bisphenols negatively impact the female germline causing oocyte defects with dramatic consequences such as aneuploïdy. We also reveal that bisphenols effects are mediated by DNA oxidation. Numerous toxicological studies have linked prophase I alterations induced by pollutants, aneuploidy and folliculogenesis defect and, here, we demonstrated the central role of oxidative DNA damages in these ovarian reproductive failures (Cuenca et al., 2020; Gely-Pernot et al., 2017; P. A. Hunt et al., 2012; Susiarjo et al., 2013). This opens new research avenues considering DNA oxidation in the developing germline as the cause of adult reproductive defects. Such mechanism remains to be investigated in the human germline and could also be invoked for numerous pollutants, either considered or not as endocrine disruptors, whose reprotoxic potential has poorly been studied despite a strong oxidative potential.

## Experimental Procedures

### Animals and gonads collection

All animal care protocols and experiments were reviewed and approved by the ethics committee of CETEA–CEA DSV (France, APAFIS 18515-2019011615312443v1) and followed the guidelines for the care and use of laboratory animals of the French Ministry of Agriculture. Mice were maintained in used polypropylene cage in standard and controlled photoperiod conditions (lights on from 8 a.m. to 8 p.m.) and had free access to tap water incubated in glass bottle and food. Males and females were caged together overnight (one male for 2 females), and the presence of vaginal plug was examined the following morning. The day following overnight mating is counted as 0.5 day post conception (dpc). All mice were killed by cervical dislocation at 11.5 dpc, 12.5 dpc, 14.5 dpc, 18.5 dpc and 8 dpp or at 3 months after birth. For pregnant mice, fetuses were removed after euthanasia from uterine horns before gonad isolation under a binocular microscope. The mice used in this study were NMRI mice (Naval Maritime Research Institute) and OGG1 deficient C57/Bl6 mice obtained from the production colony at our laboratory (Klungland et al., 2002).

### Exposition protocol

The exposition protocol of this study is shown in Figure 2C and Supplementary Figure 2. BADGE (CAS number 1675-54-3, Sigma D3415) and BPAF (CAS number 1478-61-1, Sigma 257591) were dissolved in absolute ethanol (puriss, ≥99.8%). Final concentration of 10 µM (diluted in ethanol 0.1%) were provided in drinking water to isolated pregnant females from 10.5 dpc to 14.5 dpc for short-term exposure (Supplementary Figure 2) or to 18.5 dpc for long-term exposure (Figure 2C). The control group was given drinking water added with 0.1% ethanol. For transcriptomic analyses, pregnant mice were euthanasied at 11.5 dpc and 13.5 dpc to collect gonad for germ cell sorting. As one adult mouse drinks 150 ml per kg body weight and per day, the evaluated daily intake of BPAF and BADGE by treated mice is ∼500 µg/kg/day. Internal BPAF concentration was evalueted. Total BPAF was measured by gas chromatography coupled to tandem mass spectrometry (GC-MS/MS) in the plasma of 18.5 dpc pregnant mice as previoulsy described (Eladak et al., 2018) The mean ± sem values in the plasma of BPAF group were 5.96 ± 0.78 ng/ml (n=4). As a comparison, he ‘European Food Safety Authority (EFSA) stated that the NOAELs for BADGE and BPAF is 15 and 30 mg/kg/d respectively https://www.anses.fr/fr/system/files/CHIM2009sa0331Ra-1.pdf. N-acetylcystein (NAC; Sigma A9165, 4 mM) and potassium bromate (KBrO3; Merck 1.04212.0250 0.15 mM) were also provided in water drink from day 10.5 dpc to the end of experiment.

### BrdU incorporation and detection in fetal ovaries

For BrdU incorporation, 14.5 dpc ovaries were cultured in hanging drops with BrdU (1%). After three hours, ovaries were fixed in 4% paraformaldehyde and processed for histology.

### Histology and immunofluorescence on ovarian sections

Protocols used have been described previously (Abby et al., 2016, Poulain et al., 2014). Briefly, fetal and adult ovaries were fixed overnight in Bouin’s fluid. After being dehydrated and embedded in paraffin, gonads were cut into 5-μm-thick sections. Sections were then mounted on slides for haematoxilin and eosin coloration. Germ cells on each section were identified on the basis of their histological features, as previously described and oogonia were identified as small cells with high nucleocytoplasmic ratio and the presence of prominent nucleoli. Meiotic cells displayed markedly condensed chromatin, forming distinct fine threads with a beaded appearance at the leptotene stage, and a characteristic criss-cross of coiled chromosome threads at the zygotene stage, while oocytes reaching the diplotene stage (naked or enclosed in follicle) had an increased size with the reformation of a single nucleolus. The Histolab analysis software (Microvision Instruments, Evry, France) was used for counting. Immunostaining were performed on sections from fetal gonads (12.5 and 14.5 dpc) fixed overnight in 4% paraformaldehyde (PFA). Sections were submitted to antigen retrieval with citrate buffer (pH 6) and then blocked in Normal Horse Serum (Impress HRP Reagent Kit MP-7402) or 2% gelatin, 0.05% tween, 0.2% BSA for one hour before adding antibodies. Primary antibodies used in this study were as follows: monoclonal mouse anti-5mC (abcam ab10805, 1:100), monoclonal rabbit anti-Stra8 (abcam ab49602; 1:1000), monoclonal rat anti-TRA98 (abcam ab82527, 1:500), and monoclonal mouse anti-SYCP3 (abcam ab97672, 1:500) antibodies were used. Specific donkey secondary antibodies conjugated with either Alexa Fluor 488 or 594 (1:500). BrdU detection in histological sections was performed with the Cell Proliferation kit (RPN20, GE Healthcare). Slides were mounted with Vectashield medium. Images acquisition was accomplished with a Leica DM5500 B epifluorescence microscope (Leica Microsystems) equipped with a CoolSNAP HQ2camera (Photometrics) and ImageJ Software. Images were analyzed with the Image J software.

### Immunofluorescence on chromosome spreads

Chromosome spreads were prepared using fetal gonads (12.5 and 18.5 dpc). Fetal ovaries were lacerated on precleaned/ready-to-use superfrost slides in 1X PBS, then they were supplemented with 0.2% sucrose before adding 1% paraformaldehyde/1% Triton. Slides were incubated for 1 h at room temperature in a humid chamber and then dried under a hood and then washed two times for 10 min with 0.4% H2O/Photoflow (Sigma-Aldrich). Slides were stained immediately or dried and stored at −20 °C. Spreaded 12.5 dpc oogonia were rehydrated in PBS, denaturated with 2N HCl for 45 minutes and then neutralized with a 50 mM Tris-HCl (pH8.8) solution. Slides were washed three times in PBS-0.1% Triton and then incubated in blocking solution (PBS with 0.1% triton and 5% donkey serum) before adding antibodies. Monoclonal mouse anti-8-OHdG (Eurobio MOG-020P, 1:200) and monoclonal rat anti-TRA98 antibodies were diluted in blocking solution and incubated overnight. Spreaded 18.5 dpc oocytes were blocked in blocking solution before adding antibodies. Monoclonal mouse anti-MLH1 (BD Bio sciences, G16815, 1:50) and monoclonal rabbit anti-SYCP3 (Novus, NB300232, 1:500) antibodies were diluted in blocking solution and incubated overnight. After 3 washes, donkey secondary antibodies conjugated with either Alexa Fluor 488 or 594 (1:500) were added. Metaphase I and II oocytes were obtained with adult ovaries (3 months old females). Ovaries were lacerated in M2 medium (Sigma M7167) and oocytes at germinal vesicle stage were collected and cultured for five hours (for metaphase I) or sixteen hours (for metaphase II) in M16 medium, covered with mineral oil. Oocytes at Germinal Vesicle Breakdown (GVBD) stage (for metaphase I) or with the first polar globule expulsion (for metaphase II) were transferred into a drop of Tyrode solution for zona pellucida removal. They were spread on Teflon printed diagnostic slides in water with 3mM DTT, 0.15% Triton and 0.6% paraformaldehyde. Slides were then dried and stored at −20 °C. Slides were washed three times in PBS and then incubated in blocking solution (PBS with 2% gelatin, 0.05% tween, 0.2% BSA) before adding monoclonal human anti-CREST (1:1000) antibody overnight. After 3 washes, goat anti-human secondary antibodies conjugated with Alexa Fluor 488 (1:500) were added. Slides were mounted with Vectashield with or without DAPI medium. Images were processed and specific structures were quantified with the ImageJ software (Cell Counter plugin).

### Germ cell isolation

11.5 dpc and 13.5 dpc germ cell isolation using SSEA-1 antigen was performed as previously described (Guerquin et al, 2015). Briefly, fetal ovaries were dissociated using trypsin solution. Cells are incubated with anti-SSEA (1/5, anti SSEA1 monoclonal antibody DSHB) and then incubated with a microbead-linked donkey anti-mouse IgM antibody (Miltenyi Biotec). Then, cells were applied onto an MS+ column (Miltenyi Biotec) and the positive fraction was flushed after the removal of the magnet.

### Affymetrix sample preparation

After isolation, cells were centrifuged and resuspended in RNeasy lysis buffer. RNA was extracted using Qiagen Rneasy miniKit as recommended by the manufacturer. Total RNA concentration and RNA integrity was monitored by electrophoresis (Agilent Bioanalyzer; RNA 6000 Pico Assay). Three pools of cells from 20-30 fetus (5 independent exposures) were used for differential expression analyses. Gene expression analysis was conducted using Mouse Clariom S array (Thermo Fisher) at 11.5 dpc and a Mouse Clariom D array at 13.5 dpc (Thermo Fisher). 500 µg of total RNA were processed according to the manufacturer. Raw data were generated and controlled with Expression console (Affymetrix) at the Genomic’s platform (Institut Cochin, Paris) for 11.5 dpc analyses and at the CEA for 13.5 dpc analyses.

### Microarray analysis

For gene expression analyses (11.5 dpc or 13.5 dpc), R oligo package 1.42.0 (Carvalho & Irizarry, 2010) was used for probeset annotation with expression summarized at the transcript level and robust multi-array averaging-normalized. Differential expression testing was conducted by ANOVA after linear model fitting of expression intensities with the limma R package. Microarray expression values are represented as log(2) normalized intensities. Differentially expressed genes (DEG) was filtered with a logFC ≥ 0.5 and a *pvalue* < 0.05. For 13.5 dpc RNA splicing analyses, the expression was summarized at the exon level. Samples were analyzed using default parameters. An alternative splicing event was defined as a differentially expressed exon with at least a 1 Log fold change in expression, at a *pvalue* of < 0.05.Downstream analyses were performed with R version 3.5.0 on a CentOS Linux 7 system (64-bit).Enrichment analyses were performed using ClusterProfiler, a R package for comparing biological themes among gene clusters (G. Yu et al., 2012).

### Splicing variant detection by quantitative PCR

Total RNA from isolated germ cells was extracted using the RNeasy minikit (QIAGEN, Valencia, CA, USA). cDNA was obtained by reverse transcription using the high capacity kit (Applied Biosystems, Foster City, CA, USA) according to the manufacturer’s instructions. Set of primers specific to the truncated/spliced region and reference primers located to the core/unspliced of the transcript were used to quantify the region of splicing using 2 delta delta CT methods.

### Statistical analysis

All data are presented as means ± s.e.m. Statistical analyses were performed using Graphpad software and the R version 3.5.0. All individual biological replicates were randomly sampled from 3 independent exposures (for bisphenols exposure). The statistical significance in the difference between control and bisphenols-treated data were evaluated using the non-parametric test Mann-Whitney. Statistical significance was set as p<0.05.

### Data and Code Availability

Datasets are available at the European Bioinformatics Institute under accession n° (E-MTAB-9344). The authors declare that the data supporting the findings of this study and custom code generated for the manuscript are available from the Lead Contact upon request.

## Acknowledgments

We are grateful to Katja Wassmann, Damien Cladière and Eulalie Buffin for discussion and suggestions and technical help for metaphase I and II preparation. We also thank A. Gouret and A. Leliard for her skillful secretarial assistance. We also thank the team within the animal housing facility at the iRCM. For affymetrix data, we thank Sébastien Jacques and Angeline Duché from Genom’ic (institut Cochin). For BPAF concentration in serum samples, we thank jean-Philippe Antignac from LABERCA (Oniris, INRAE, Nantes). This research was supported by the ADEME (French Environment & Energy Management Agency), ANSES (French Agency for Food, Environmental and Occupational Health & Safety) Université de Paris and INSERM. S.A. were supported by a fellowship from the Ministère de l’Enseignement et Recherche.

## Author Contributions

Conceptualization: M.J.G, G.L., V.R.F, A.C, J.P.R, E.M and R.H; Methodology: S.A., M.J.G, A.C, J.P.R, E.M; Validation: S.A., D.M., M.W and S.M.; Formal Analysis: M.J.G and S.A. Investigation: S.A., D.M., M.W and S.M; Resources: A.C. and J.P.R; Data Curation: M.J.G.; Writing –Original Draft: S.A. and M.J.G.; Writing –Review & Editing: S.A., G.L., V.R.F, A.C, J.P.R, E.M and R.H. Visualization: S.A. and M.J.G. Supervision: M.J.G, V.R.F and G.L. Project Administration: M.J.G, V.R.F and G.L. Funding Acquisition: M.J.G, V.R.F and G.L.

---

**Supplementary Figure 1:**
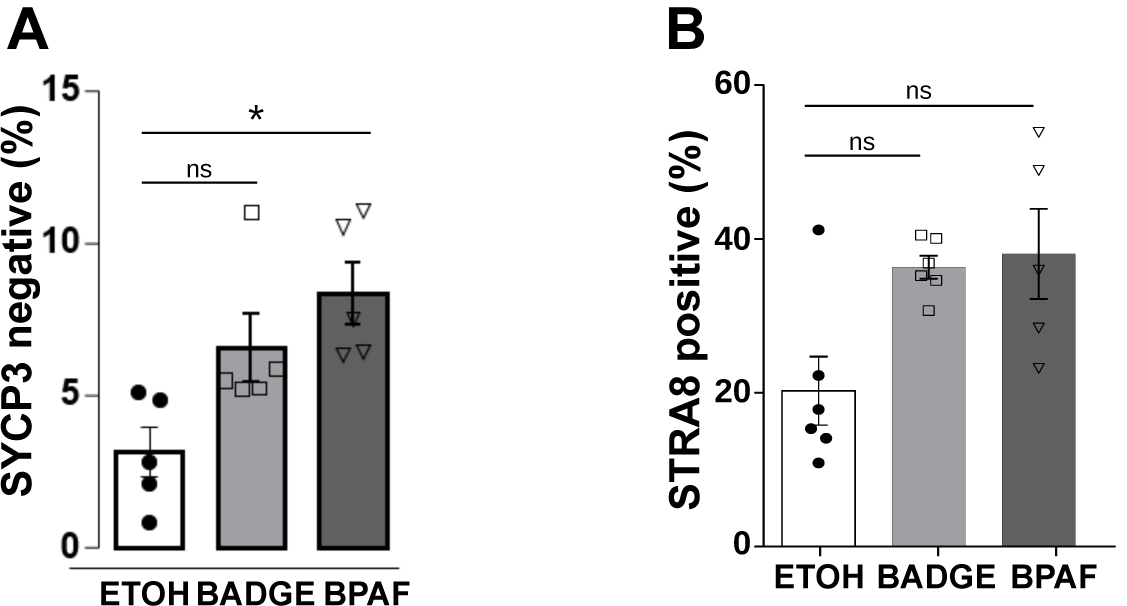
(A) Percentage of oogonia (SYCP3-/ TRA98 positive). Error bars show mean ± s.e.m. n=5 mice from independent exposures; * *p <*0.05 (Mann-Whitney’s test).(B) Percentage of germ cells expressing STRA8 in 14.5 dpc female gonads. n=5 mice from independent exposures; p= 0.08 (BADGE), p= 0.07(BPAF) (Mann-Whitney’s test).

**Supplementary Figure 2:**
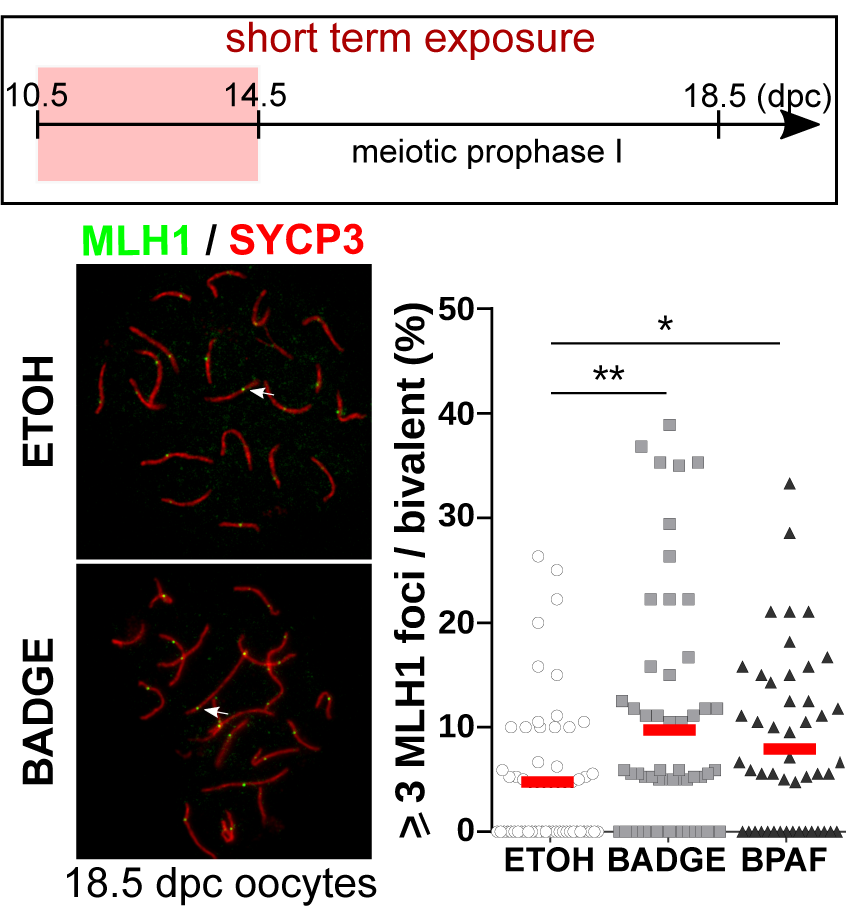
Ovaries from fetuses exposed to vehicle (ETOH) or bisphenols (BADGE and BPAF) from 10.5 dpc to 14.5 dpc were used for the MLH1 quantification (short exposure). Left panel: Representative photomicrographs of pachytene cells from vehicle (ETOH) and BADGE treated ovaries. White arrow shows a synaptonemal complexe with 3 MLH1 foci. Right panel: Percentage of synaptonemal complexes with 3 or more MLH1 foci. n= 62 (ETOH), 55 (BADGE) and 55 (BPAF) oocytes from 3 independent exposures. Mean (red bar), * p < 0.05, ** p < 0.01 (Mann-Whitney’s test).

**Supplementary Figure 3.**
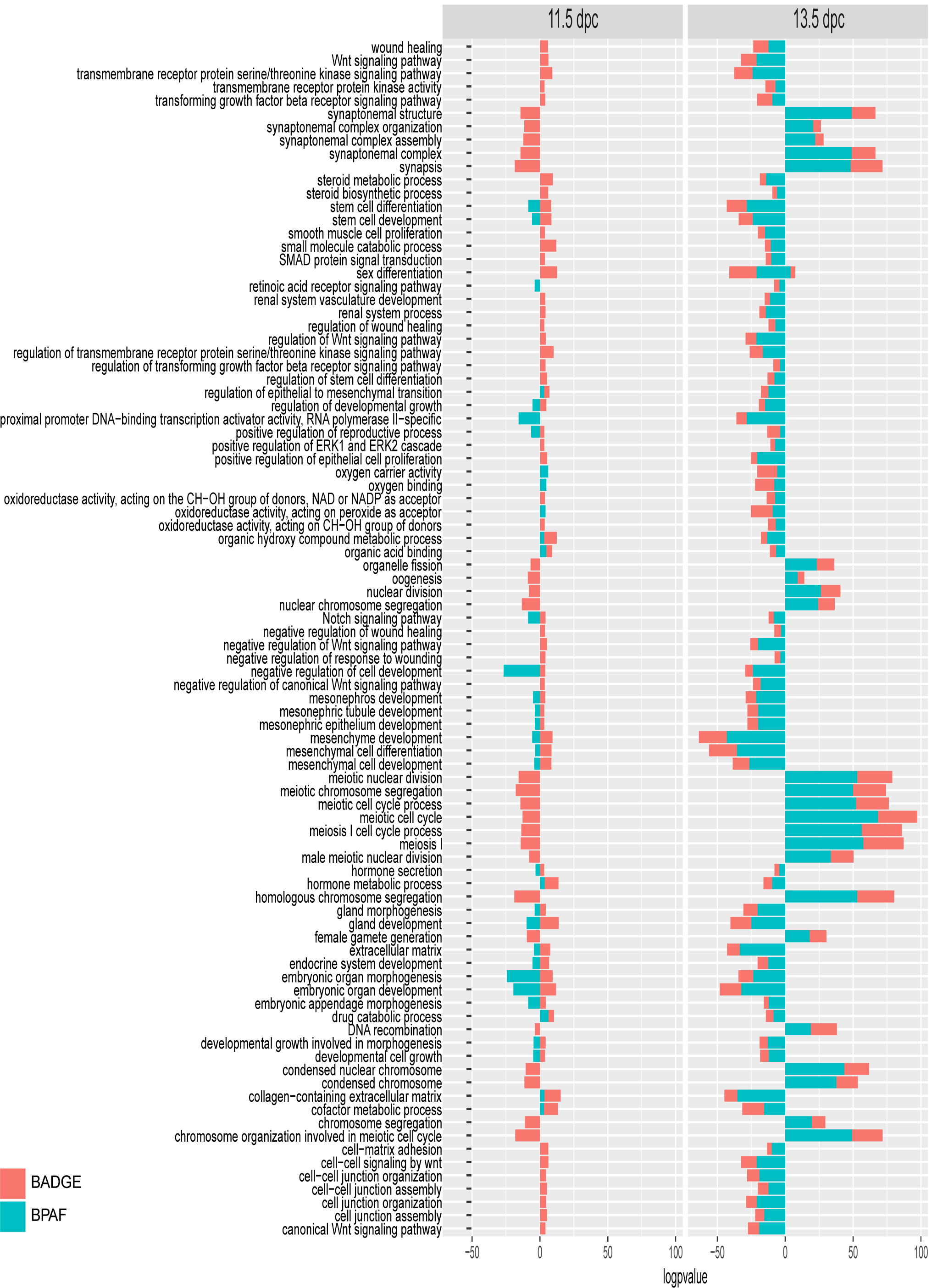
Enrichment of Gene Ontology (biological process, molecular function and cellular component) of DEGs after bisphenols exposure in 11.5 and 13.5 dpc. Ontologies associated to down-regulated genes is presented as negative values of *pvalues*.

**Supplementary Figure 4:**
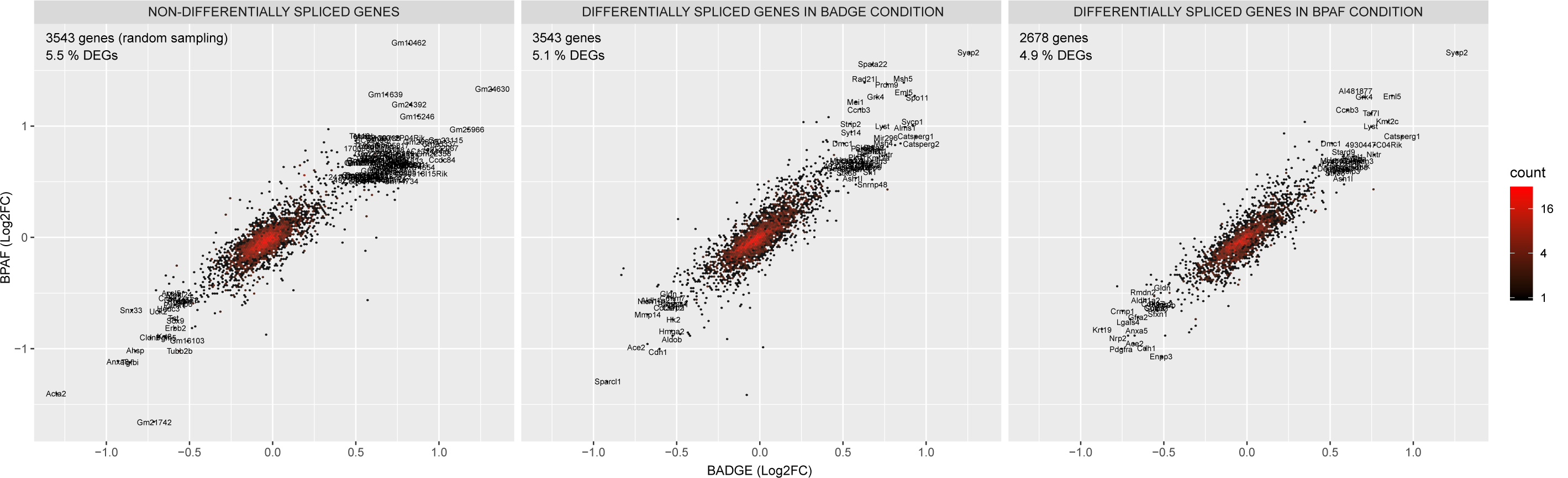
2d density plot between BADGE and BPAF condition of the differential expression (Log2FC) of differentially spliced genes in BADGE or BPAF condition. The plot area is divided in 200 hexagons and the color of the hexagon correspond to number of genes inside the hexagon. A simple random sampling of 3543 non differentially spliced genes (maximal number of differentially spliced genes) was used to represent transcriptionnal behavior after BADGE or BPAF exposure of non differentially spliced genes.

**Supplementary Figure 5.**
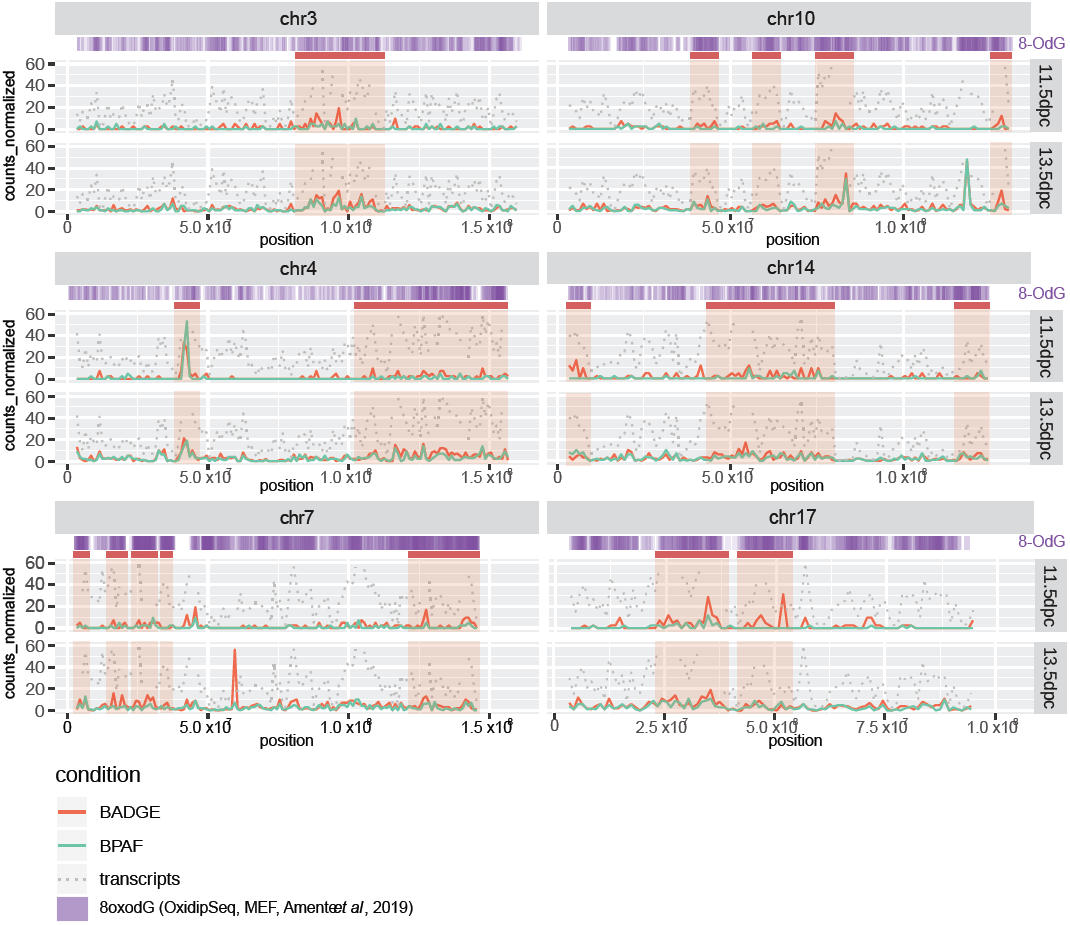
Genome mapping on selected chromosomes (Chr 3, 4, 7, 10, 14, 17) of the DEGs in BPAF (red line) and BADGE (green line) conditions. The sum of the filtered genes per megabase (1000000 bases) was reported according to their location on the chomosome. Grey dotted line represent the sum of the total genes referenced in the murine genome. Purple line correspond to the mapping of the OxiDIP-Seq signal r profiles retrieved in the Amente et al publication (Amente et al., 2019).

**Supplementary Figure 6:**
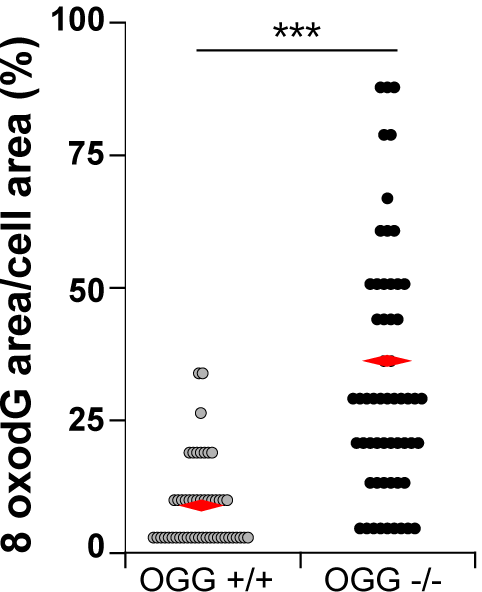
Quantification of the total area of 8OdG normalized by the nucleus area in 12.5 dpc OGG +/+ and -/-oogonia .n= 51 (OGG^+/+^), n=58 (OGG^-/-^). *** p<0.001 (Mann-Whitney’s test).

